# Likelihood Ratios Given Activity-Level Propositions for DNA Transfer Evidence: Theoretical Foundations of the HaloGen Framework (Part I)

**DOI:** 10.64898/2026.02.03.702484

**Authors:** Peter Gill, Øyvind Bleka

## Abstract

The interpretation of trace DNA evidence at activity level requires explicit modelling of transfer, persistence, and the possibility that a relevant actor leaves no detectable DNA. We present the theoretical foundations of HaloGen, an open-source hierarchical Bayesian framework for evaluating DNA quantities under competing activity-level propositions.

The propositions are framed in terms of alleged activities or actions; direct and secondary transfer are treated as transfer mechanisms through which those activities may give rise to the observed DNA. HaloGen accounts for zero transfer, multiple contributors, specified unknown contributors, unobserved actors, and multiple stains. Evidence is evaluated using an exhaustive-propositions likelihood-ratio framework that combines information across contributors and stains while propagating uncertainty in transfer and detection. Observed DNA quantities and non-detects are handled within a single probabilistic framework: detected quantities are evaluated using conditional-on-detection densities, whereas the probability that a relevant actor leaves no detectable DNA is represented by an empirically constrained fail-rate parameter.

The framework yields transparent and stable behaviour: informative DNA quantities can support propositions involving direct transfer, while low-information or no-detect situations are neutral or defence-conservative. The empirical-clamped fail-rate policy prevents spurious inflation of likelihood ratios when non-detection of a relevant actor is plausible. This paper establishes the theoretical basis of HaloGen; a companion paper addresses validation and applied casework examples.

## 1. Introduction

Interpreting trace DNA quantities recovered from touched items remains one of the most challenging tasks in activity-level evaluation. Activity-level propositions should be framed in terms of alleged human activities or actions, such as handling, prior social contact, assault, or legitimate use. Direct, secondary, or background transfer are not themselves the propositions; rather, they are biological transfer mechanisms through which the activities described in the propositions may give rise to the observed DNA. A robust assessment therefore requires probabilistic models that describe how DNA quantities arise under the transfer pathways implied by the competing propositions [1]. For any given activity, these probabilities can vary substantially between laboratories due to differences in sampling, extraction, and amplification procedures [2, 3]. The likelihood ratio (LR), formally equivalent to a *Bayes Factor* (BF), provides a logically coherent measure of the relative support for competing propositions [4]. Furthermore, hierarchical Bayesian models (e.g., random-effects models) provide a natural way to represent inter- and intra-laboratory variability [5, 6] However, such statistical structure alone is not sufficient for activity-level interpretation, which also requires explicit quantitative models of the biological mechanisms of transfer and their associated uncertainty, including the possibility of unobserved relevant actors or undetected DNA [7].

A central consideration at this level of interpretation is the presence of background DNA. While sometimes treated as “nuisance” under the prosecution proposition (*H*_*p*_), background DNA can represent genuine alternative contributors under the defence proposition (*H*_*d*_). Unknown contributor probabilities must be explicitly modelled within the LR framework; otherwise, the analysis risks overstating the weight of evidence for the person of interest. Experimental findings from ReAct Experiment 3 [2] showed that, under the particular post-handshake conditions examined in that study, background or indirectly transferred DNA quantities could overlap with secondary-transfer distributions. This observation should not be read as a general claim about all background or secondary-transfer scenarios, since different activities, contact durations, substrates, contributors and sampling strategies may produce substantially different quantity distributions. Before any activity-level assessment can be made, the DNA quantity attributable to each contributor must be estimated. This typically involves two stages: (i) quantifying the total DNA using qPCR, and (ii) estimating mixture proportions to apportion this total among contributors. Probabilistic genotyping systems such as *EuroForMix* [8] are widely used at the sub-source level, modelling electropherogram peak heights to compute sub-source LRs. Combining these mixture proportions with the total DNA yield provides an estimate of the absolute DNA quantity attributable to each contributor. While this quantitative information is essential, it addresses only the *sub-source* level; the central question of *activity-level relevance*, what those quantities imply about the mechanisms of transfer, is a different consideration.

Previous efforts have developed quantitative probabilistic models for this purpose. Bright et al. [9] introduced log normal distributions to model DNA quantities, and others have explored extensions using alternative likelihood formulations [10, 11]. However, when these approaches are applied to results given activity propositions, they typically assume simple experimental conditions: one stain, one contributor, and fixed background behaviour. In complex casework involving multiple known and unknown contributors in several stains, a more general approach is required, one that can systematically assess all logically possible combinations of the presence and absence of the contributor, without relying on fixed assumptions about the probability or quantity of background

The *exhaustive approach* to forensic evidence interpretation provides precisely such a framework [12, 13, 14]. Under this approach, every admissible subset of known/ unknown contributors (or “elemental hypothesis”) is enumerated, and the likelihood for each is computed. This exhaustive set forms a closed and mutually exclusive partition of all possible evidential explanations. The overall LR is then derived by summing over the likelihoods of all elemental hypotheses consistent with each top-level proposition (*H*_*p*_ and *H*_*d*_).

Building on this foundation, the present paper introduces the **HaloGen** (*Hierarchical Bayesian Activity-Level Orchestrator: next Generation*) framework - a generalised, hierarchical Bayesian implementation of the exhaustive approach for activity-level DNA transfer analysis. HaloGen extends existing quantitative models as follows:

1. **Contributor-specific evaluation:** Each specified contributor, including each unknown, is treated as a distinct entity rather than aggregating all non-POI DNA into a single “background” term.
2. **Generalisation to multiple relevant actors:** The exhaustive-hypothesis method is extended to scenarios involving multiple possible actors, a frequent feature of complex casework.
3. **Explicit modelling of unobserved actors:** HaloGen introduces a fail-rate probability (*F*_0_) to represent potential actors who left no detectable trace.

This first paper in a two-part series presents the theoretical foundations of the HaloGen model. This is summarised in Section 2; section 2.4 details the extension of the exhaustive framework, previously applied at sub-source level [13, 14], to the activity-level domain. Section 2.3 introduces the multistain extension, reflecting common casework practice. Section 3.2.3 examines how the inclusion of unknown contributors influences the likelihood ratio for the person of interest. Section 3.3 introduces the anchor principle, a conceptual safeguard ensuring meaningful and balanced interpretation, and Section 3.5 outlines prospective extensions, including models for individual shedder variability.

Computational Bayesian estimation methods are detailed in S3. A companion paper (Part 2) will present empirical validation, software implementation, casework applications showing results from analyses from 20 different laboratories.

## 2. Methods

This section outlines the statistical framework implemented in HaloGen for evaluating DNA transfer evidence at the activity level https://github.com/peterdgill/HaloGen_Bayes. HaloGen quantifies evidential strength using a Bayes factor (BF), reported on the likelihood-ratio (LR) scale. For each posterior draw of the transfer-model parameters, the case-level likelihood ratio is computed under the competing propositions. The reported point estimate is the median of this posterior Monte Carlo distribution of LRs, accompanied where appropriate by uncertainty intervals. The reported LR therefore reflects both the expected behaviour of the DNA transfer process and uncertainty in the estimated transfer parameters.

The workflow proceeds in three logical stages:

1. **Experimental Modelling:** Defining the probability distributions for DNA transfer and estimating their parameters from laboratory data.
2. **Case-Level Transition:** Converting these experimental parameters into conditional likelihoods (*t*_*c*_, *s*_*c*_) and non-detect probabilities (*F*_0_) applicable to casework.
3. **LR Construction:** Combining these probabilities into an exhaustive likelihood ratio that accounts for multiple contributors and stains.

A glossary of notation is provided in S1.

### 2.1. Experimental Transfer Model

HaloGen models laboratory transfer data using a zero–augmented and left–censored Lognormal mixture model. Each experimental observation arises from one of two mechanisms:

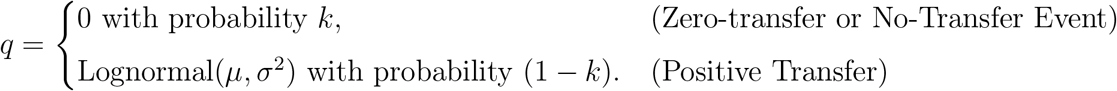

Either a zero-transfer or no-transfer event, with probability *k*, in which no measurable DNA is transferred from the donor, or a positive transfer, with probability 1 − *k*, in which the amount of DNA transferred follows a lognormal distribution. Positive transfer may nevertheless fall below the detection limit (DL) and therefore be recorded as a non-detect (Fig. 1). The model assumes that, conditional on the occurrence of positive transfer, the magnitude of the positive amount is governed by the lognormal component and is independent of the zero-transfer mechanism represented by *k*. This is a simplifying distributional assumption that can be replaced if future data support a different dependence structure.

**Figure 1:**
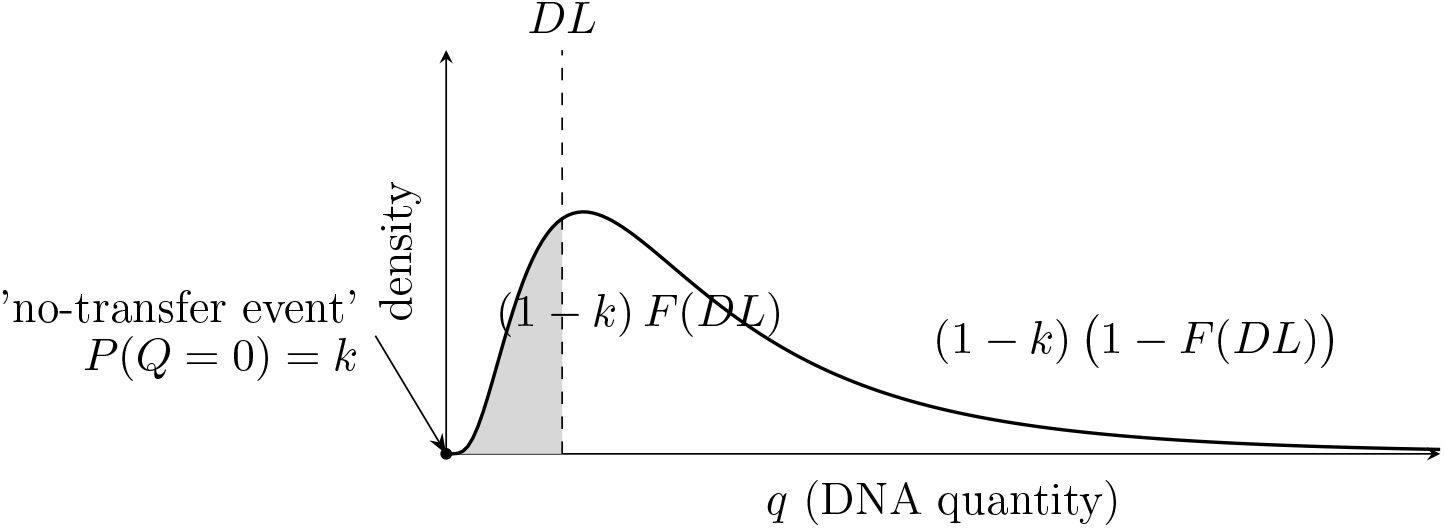
Zero-inflated, censored Lognormal representation…

The Lognormal component describes the distribution of *positive* DNA amounts. However, a laboratory detection limit DL causes censoring. Here, *F* (DL) represents the Lognormal CDF evaluated at the detection limit. Physically, this is the probability that a positive DNA amount falls into the “invisible” range below the detection threshold.

Consequently, a recorded non–detect can arise from two sources:

1. A true ‘no-transfer event’ (probability *k*).
2. A positive transfer that is hidden below the limit (so the combined probability is (1 − *k*)*F* (DL)).

The total probability of a non–detect is the sum of these distinct possibilities:

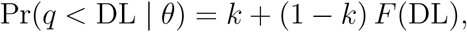

where *θ* = (*µ, σ, k*) denotes the model parameters.

Conversely, the probability of observing a positive detect is the weight of the positive mechanism multiplied by the size of the visible area of the probability density:

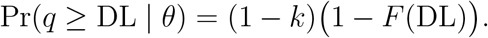

For Bayesian parameter estimation, we define the likelihood function *L*(*θ*). This requires different handling for the discrete non–detects versus the continuous detects.

For a **non–detect**, where contributor DNA cannot be observed, the likelihood contribution is the probability mass of this event:

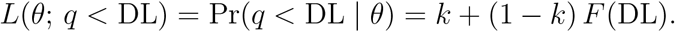

For a **detect** (*q* ≥ DL), we observe a specific continuous value. Conditional on a positive transfer having occurred, the likelihood contribution is proportional to the lognormal density *f*_LN_(*q*), weighted by the probability of the positive mechanism (1 − *k*):

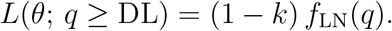

#### 2.1.1. Difference between detection limit and analytical threshold

It is important to distinguish between the analytical threshold (AT), which operates at the sub-source level, and the detection limit (DL) that governs the activity-level model. The AT is primarily a noise-filtering device and is typically set deliberately high to avoid spurious allele calls. Activity-level TPPR studies often involve limited sample sizes, sometimes fewer than 20 usable observations for a given activity, laboratory, substrate or time interval. For that reason, it is important not to discard low-level measurements unnecessarily by imposing a high activity-level threshold. The detection-limit modelling boundary used here is intended to retain the maximum amount of relevant quantitative information while still respecting the laboratory’s ability to distinguish detected from non-detected quantities.

In our setting the laboratory experiments are performed under controlled conditions where the ground truth is known: specifically, which transfers are direct, which are secondary, and the corresponding quantitative outcomes. This allows us to model all parameters of the mixture distribution (including the zero-mass, log-normal positive component, and DL behaviour) directly from these data rather than relying on indirect inference from casework samples. Using the R package fitdistrplus(), we assessed how well a log-normal distribution fit the positive (above-DL) values under a range of candidate detection limits. A DL of 0.001 ng gave the best result: it retained the largest amount of usable data while still maintaining a good log-normal fit.

To assess robustness to the DL modelling boundary, we also performed a detection-limit sensitivity analysis (Supplement S2.5), recomputing likelihoods and case LRs under DL/2 and 2·DL.

### 2.2. Parameter Estimation from Experimental Data

The parameters (*µ, σ, k*) of the censored Lognormal mixture model are estimated separately for direct and secondary transfer using Bayesian inference. HaloGen uses Markov Chain Monte Carlo (MCMC) sampling (implemented in Stan [15]) to obtain posterior draws of the model parameters:

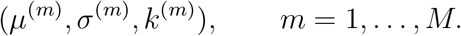

The resulting posterior samples form the basis for all downstream calculations. We write *t*_*c*_(*q*) and *s*_*c*_(*q*) for the case-level likelihood contributions for a detected contributor quantity *q* under the direct-transfer and secondary-transfer pathways, respectively. These quantities are evaluated for each posterior draw. Their formal conditional-on-detection definitions are given in Section 2.3. For any observed quantity *q*, the per-person transfer likelihoods used in the case-level analysis are obtained by evaluating:

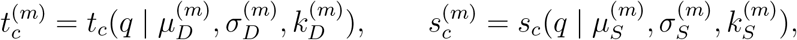

for each posterior draw *m*. HaloGen therefore propagates uncertainty in the experimental model directly into the final activity-level LR by using samples from the posterior distribution of (*µ, σ, k*), rather than relying on point estimates. Full details of the MCMC implementation, diagnostics, and convergence checks are provided in Supplement S3.1.

HaloGen provides three model variants for estimating these parameters, offering different ways to balance general knowledge with lab-specific data:

1. ***Group* Model:** This model learns general trends by pooling data across many different laboratories. It provides robust, generalised estimates of transfer parameters. This is useful when the user’s specific laboratory has limited data, but the resulting parameters may be too generic if the lab’s conditions differ significantly from the global average.
2. ***Lab–Vague* Model:** This model estimates parameters using *only* the data from the specific laboratory, with minimal external assumptions (using “uninformative” or “vague” priors). While this reflects the unique characteristics of that lab, it is susceptible to instability. If the lab’s dataset is small or unrepresentative, this model can yield large variances (high uncertainty) or, conversely, confident but erroneous conclusions driven by chance outliers in the small sample.
3. ***Lab–Bayes* Model (Gold Standard):** This is the recommended model for reporting. It provides estimates for a specific lab by combining the strengths of the other two. It employs a hierarchical approach: it starts with the general trends learned by the *Group* model (using them as *informative priors*) and then refines these estimates based on the specific data from the individual lab. This stabilizes the estimates, preventing the overfitting issues of the *Lab–Vague* model while ensuring the results remain specific to the local laboratory conditions.

The resulting posterior samples form the basis for all downstream calculations. HaloGen propagates uncertainty in the experimental model directly into the final case-level activity-level LR by using posterior samples to construct a Monte Carlo representation of the case-level likelihood ratio (see Supplement S2.4) .

### 2.3. From Experimental Distribution to Case-Level Modelling

The experimental model described above is a generative model for the quantity *Q* that may be recovered from a transfer experiment. It assigns probability both to detectable quantities and to non-detects. A non-detect may arise either because no DNA was transferred, represented by *k*, or because a positive transfer occurred but the quantity fell below the detection limit DL. Thus, under model parameters *θ* = (*µ, σ, k*),

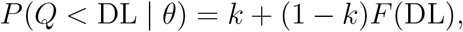

where *F* (DL) is the Lognormal distribution function evaluated at the detection limit. This generative probability is retained in HaloGen for events in which a contributor, in particular an actor, is not observed.

Case-level interpretation involves an additional conditioning step. When a contributor has been detected in a case stain and an estimated quantity *q* ≥ DL is available, the relevant question is no longer whether a detect occurred. Detection has already occurred. The relevant likelihood contribution is therefore the density of the observed quantity conditional on detection:

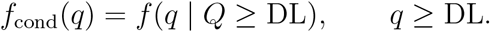

Under the zero-augmented Lognormal model, the positive-transfer component has weight (1 − *k*) and Lognormal density *f*_LN_(*q*). The unconditional probability of a detect is

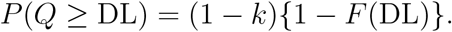

Hence, for a detected quantity,

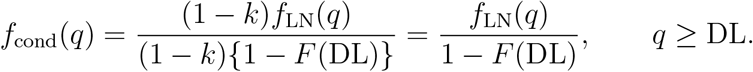

The factor (1 − *k*) cancels because we condition on the event that a detectable quantity exists. This conditional density integrates to one over (DL, ∞) and describes the distributional shape of the observed DNA quantity, given that it is detectable.

Accordingly, for a detected case quantity, HaloGen uses conditional case-level densities for both the direct and secondary transfer mechanisms:

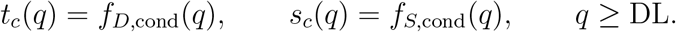

Here *t*_*c*_(*q*) and *s*_*c*_(*q*) are both conditional-on-detection case-level densities. The subscript *D* denotes the fitted direct-transfer distribution and the subscript *S* denotes the fitted secondary-transfer distribution. Thus, *t*_*c*_(*q*) is used when contributor *c* is assigned to the direct-transfer role under an elemental hypothesis, whereas *s*_*c*_(*q*) is used when contributor *c* is assigned to the secondary-transfer role. These assignments refer to roles within the activity-level proposition; they do not imply uncertainty about the contributor’s source-level identity.

For a contributor who is not detected, no continuous quantity is available and no conditional density term is evaluated. Instead, the relevant contribution is a probability mass for the non-detect event:

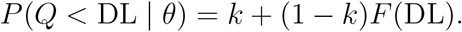

This probability-mass term is conceptually distinct from the conditional densities *t*_*c*_(*q*) and *s*_*c*_(*q*), which are defined only for detected quantities *q* ≥ DL.

In particular, when an elemental hypothesis requires a relevant actor who is not represented among the detected contributors, HaloGen uses the directtransfer non-detection probability

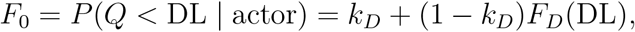

where the subscript *D* denotes the direct-transfer model. This quantity is introduced formally in Section 2.3.1. Thus, a detected contributor quantity is evaluated through *t*_*c*_(*q*) or *s*_*c*_(*q*), whereas an unobserved actor required by the hypothesis contributes through *F*_0_.

The case-level likelihood contribution for contributor *c* under elemental hypothesis *h* can therefore be written as

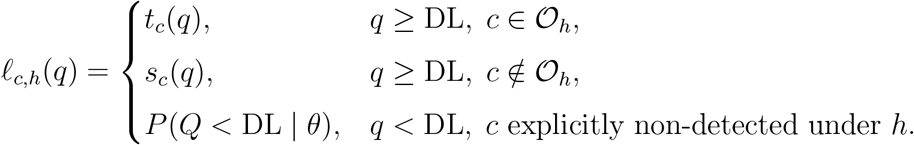

Here 𝒪_*h*_ denotes the set of contributors assigned to the direct-transfer actor role under elemental hypothesis *h*. Contributors not in 𝒪_*h*_ are assigned to the secondary-transfer role. The third line applies only where the hypothesis explicitly requires a non-detect event for contributor *c*; it is a probability-mass term, not a density.

The first two lines are conditional densities for observed quantities; the third line is a probability mass for a non-detect event. This distinction is important because likelihood ratios may combine continuous-density terms for observed contributors with probability-mass terms for non-detected or unobserved contributors.

HaloGen also allows evidence from multiple stains or items to be combined. For contributor *c*, let

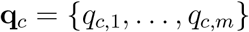

denote the case-level observations across stains, where a value *q*_*c,j*_ ≥ DL denotes a detected contributor quantity and *q*_*c,j*_ *<* DL denotes a non-detect. Conditional on the transfer parameters and on the elemental hypothesis *h*, stains are treated as independent observations. The joint contribution is therefore

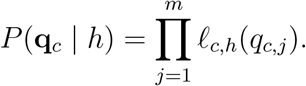

This formulation preserves the distinction between two uses of the same experimental model. The conditional densities *t*_*c*_(*q*) and *s*_*c*_(*q*) are used only for detected quantities. Non-detects are not treated as zero-valued density observations; they enter through probability mass. Most importantly, when an elemental hypothesis requires a relevant actor who left no detectable DNA, this contribution is represented by *F*_0_.

#### 2.3.1. The Actor Non-Detection Probability F_0_

We must take account of the probability that the relevant actor leaves *no* detectable DNA. Let *F*_0_ denote this case-level fail-rate:

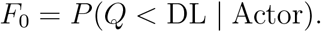

This represents the probability that the actor’s contribution is *not detected*, regardless of whether the transfer failed structurally or fell below the detection limit.

HaloGen estimates *F*_0_ using the parameters from the experimental **direct-transfer** model:

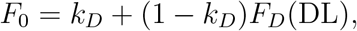

where *k*_*D*_ is the direct ‘no-transfer event’ probability and *F*_*D*_(DL) is the probability of a direct positive transfer falling into the invisible range. Thus, *F*_0_ is simply the total probability mass of the “Non-Detect” bucket derived from the Direct Transfer experiments.

This parameter is critical for evaluating the hypotheses:

- **Under** *H*_*p*_ **(Suspect is actor):** We assume the suspect is the source of the observed stain *q*_*c*_. Since *q*_*c*_ was detected, *F*_0_ does not contribute to the numerator.
- **Under** *H*_*d*_ **(Suspect is not actor):** We assume a specific unknown individual is the true actor. If the suspect’s DNA is present via secondary transfer, the true actor must have failed to leave a detectable trace. The probability of this “silent actor” event is *F*_0_.

#### 2.3.2. Ensuring Stability: The Empirical-Clamped F_0_ Policy

In an earlier iteration of the framework, the probability *F*_0_ was estimated from direct-transfer experiments using a Jeffreys-adjusted empirical Beta prior:

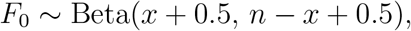

where *x* and *n* are the numbers of non–detects and total replicates in the laboratory’s direct-transfer data. This yielded realistic case-level fail rates—typically

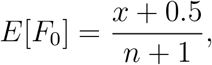

in the range 0.02–0.10 for common experimental sample sizes around *n* = 20. However, when *x* was small or zero, occasional draws from this prior could be extremely close to zero, leading to unrealistically large likelihood ratios in favour of direct transfer, because elemental hypotheses involving unobserved actors carry multiplicative factors of *F*_0_ *<* 1.

To prevent this, the *empirical-clamped* policy was introduced. It retains the same Jeffreys-smoothed empirical prior but bounds each Monte Carlo draw of *F*_0_ within a conservative interval

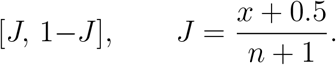

This bound is applied only to extreme outlying draws; ordinary values remain unchanged. In effect, it ensures that no case is driven by implausible tail realisations of *F*_0_ while preserving the empirical variability between laboratories. Further details are provided in Supplement S2.

### 2.4. Likelihood Ratio Construction

HaloGen uses an *exhaustive* approach to construct the likelihood ratio (LR). Every admissible detailed scenario, referred to here as an “elemental hypothesis”, is enumerated and evaluated. This general logic is related to exhaustive approaches used in multiple-person DNA mixture interpretation [14, 13, 16], but HaloGen uses the elemental hypotheses here for activity-level transfer evaluation.

Each elemental hypothesis *H*_O_ specifies a subset 𝒪 of individuals who are treated as actors under the activity-level proposition. Let 𝒦 denote the set of specified individuals, including persons of interest and other known persons, and let 𝒰 denote the set of specified unknown contributors. Let *N*_*S*_ denote the assumed number of actors required by the proposition. For an elemental hypothesis *H*_O_, the individuals in 𝒪 are assigned to the direct-transfer role, whereas those in the complement

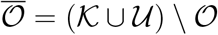

are assigned to the secondary-transfer role.

The likelihood of the evidence under an elemental hypothesis is

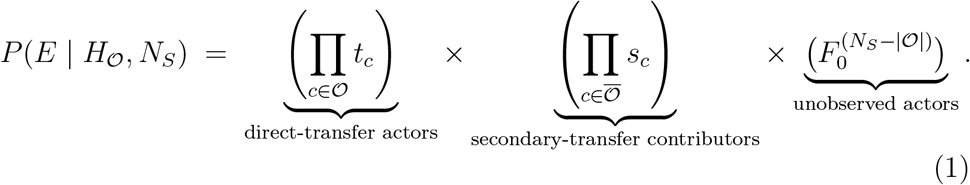

The final factor accounts for any unobserved actors needed to reach the assumed total *N*_*S*_. Such an actor is required by the proposition but is not represented among the detected contributors.

#### Elemental weighting convention

HaloGen uses an equal-elemental-weight summation convention over the closed set of admissible elemental hypotheses. This convention is not intended to assert that all elemental scenarios are equally probable in the real world. Rather, it is a transparent modelling convention for combining exhaustive elemental likelihoods without introducing proposition-dependent scaling artefacts.

Under this convention, each admissible elemental hypothesis is assigned the same common weight before summation. Because the same common weight applies to every admissible elemental hypothesis, it cancels in the LR and explicit prior-weight terms are omitted from Eq. (1). The resulting LR is driven by the total summed likelihood support for the elemental hypotheses associated with the competing propositions.

This differs from an approach in which likelihoods are normalised separately within each top-level proposition. Separate normalisation can be appropriate when explicit proposition-level prior structures are being specified, but it also means that the LR depends on how prior mass is distributed within *H*_*p*_ and *H*_*d*_. In HaloGen, the default equal-elemental summation is used as a neutral baseline when no defensible case-specific information is available to prefer one admissible actor assignment over another. If such information is available, unequal elemental weights could be introduced explicitly, but those weights would form part of the proposition structure and should be justified and explored by sensitivity analysis.

#### Effect of non-detects on the LR

Non-detects enter the exhaustive likelihood through probability-mass terms rather than through continuous densities. Their effect on the LR depends on which elemental hypotheses require one or more unobserved actors. In cases where a proposition requires an actor who left no detectable DNA, the corresponding terms include factors of *F*_0_. Because *F*_0_ *<* 1, such terms are down-weighted relative to hypotheses involving detected actors. The direction and magnitude of the overall LR effect therefore depend on the complete set of elemental hypotheses, the assumed number of actors, and the observed quantities for the detected contributors.

The LR for any specified individual is then obtained by summing Eq. (1) across all admissible elemental hypotheses in which that individual is assigned to 𝒪, and dividing by the corresponding sum across all admissible elemental hypotheses in which that individual is not assigned to 𝒪. Thus, for a specified contributor *x*,

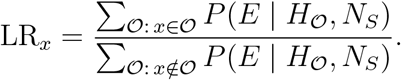

### 2.5. Worked Examples

#### 2.5.1. Case-Level Likelihoods: The One-Actor, One-Stain Example

This simple example illustrates the fundamental logic. We consider a scenario with one Person of Interest (POI) *c*, one observed contributor quantity *q* ≥ DL, and one relevant actor under the activity-level proposition, so that *N*_*S*_ = 1.

##### Prosecution proposition H_p_

Under *H*_*p*_, the POI is the relevant actor and the observed quantity is evaluated under the direct-transfer pathway. The likelihood contribution is therefore the direct conditional density:

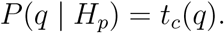

##### Defence proposition H_d_

Under *H*_*d*_, the POI is not the relevant actor. The observed POI quantity is therefore evaluated under the secondary-transfer pathway, while the relevant actor required by the defence proposition is unobserved. Two likelihood components are therefore required: the secondary-transfer density for the POI’s observed quantity, *s*_*c*_(*q*), and the non-detection probability *F*_0_ for the unobserved actor. Thus,

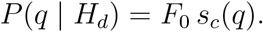

##### Resulting LR

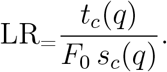

This one-stain example illustrates the component likelihood terms used by HaloGen: conditional densities for observed contributor quantities and probability-mass terms for unobserved actors. The next example illustrates how these terms are combined into a composite LR under the exhaustive hypothesis framework.

#### 2.5.2. Exhaustive LR Example: Two Known Individuals and One Specified Unknown Profile

Consider a case with three detected contributors: two known individuals (S1, S2) and one specified unknown profile (U3). We assume that the activity-level proposition requires exactly *N*_*S*_ = 2 relevant actors. Let *t*_*c*_, *s*_*c*_, and *F*_0_ denote the per-contributor case-level likelihood components defined above.

Each elemental hypothesis *H*_O_ selects a subset of 𝒪 the detected profiles and assigns those co<u>nt</u>ributors to the direct-transfer actor role. Contributors in the complement 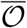 are assigned to the secondary-transfer role. If fewer than *N*_*S*_ detected contributors are assigned to the direct-transfer role, the hypothesis is supplemented by the required number of unobserved actors, each contributing a factor *F*_0_.

The likelihood is given by Eq. (1):

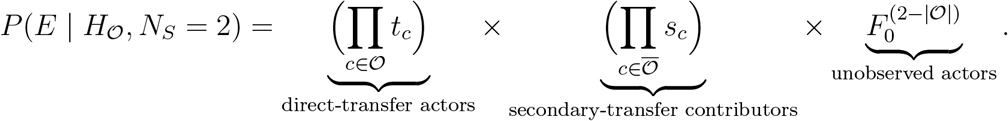

##### Examples

- **Hypothesis: S1 and S2 are direct-transfer actors**. The specified unknown profile U3 is assigned to the secondary-transfer role.

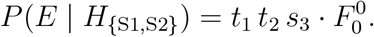
- **Hypothesis: S1 and U3 are direct-transfer actors**. S2 is assigned to the secondary-transfer role.

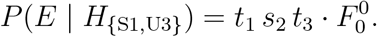
- **Hypothesis: S1 and one unobserved actor satisfy the proposition**. S2 and U3 are assigned to the secondary-transfer role.

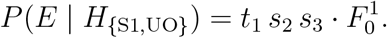
- **Hypothesis: two unobserved actors satisfy the proposition**. All detected contributors are assigned to the secondary-transfer role.

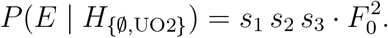

To compute LR_S1_, where S1 is the POI, we sum the elemental hypotheses in which S1 is assigned to the direct-transfer actor role in the numerator, and the elemental hypotheses in which S1 is not assigned to that role in the denominator:

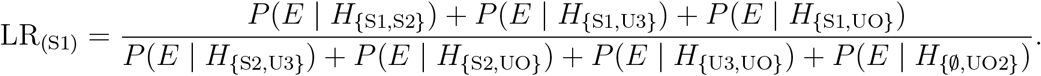

Assuming a single exchangeable prior over elemental hypotheses, the prior terms cancel in the LR.

The same exhaustive summation protocol is used to compute LR_S2_ and LR_U3_, replacing the numerator by the set of elemental hypotheses in which the contributor of interest is assigned to the direct-transfer actor role. This follows the same exhaustive likelihood-ratio logic described in [14, 13].

### 2.6. Data used in the analysis

To demonstrate the practical application of the HaloGen model, we utilised data from Experiment 3 of the ReAct study [2]. This dataset was specifically designed to investigate secondary versus direct transfer events in a controlled setting. It was collected following the recommendations of [17], except that NIST standards were not compared (since the experiments predated this recommendation).

The experiment simulated a contact scenario: a ‘simulated actor’ shook hands with a work colleague. After a delay of one hour, the actor handled a screwdriver. The screwdriver handle was subsequently analysed to determine the recovery of DNA from:

- The actor (Direct Transfer: Hand → Screwdriver).
- The Colleague (Secondary Transfer: Colleague → actor’s Hand → Screwdriver).

The dataset comprises results obtained from 20 participating laboratories. These data were analysed using the Group, *Lab–Bayes*, and *Lab– Vague* models described in Section 2.2. To provide a focused demonstration of the model’s outputs, we selected one representative laboratory, identified as `lab_1_ESS`, which demonstrated robust levels of DNA recovery. An extended analysis encompassing all participating laboratories is presented in the companion paper [18].

The use of ReAct Experiment 3 data in this paper is illustrative and does not imply that this experimental design will match every case. In casework, the expert must assess whether the available TPPR data are relevant to the disputed activities, substrate, time interval, sampling method, laboratory process and contributor population. Experimental studies rarely reproduce the case circumstances perfectly. Where the available data are only a proxy, that proxy status should be disclosed; where no scientifically defensible proxy exists, the model should not be used to offer a quantified activity-level opinion for that issue.

### 2.7. Resources

A great deal of information is available in the Supplements for those interested in details:

Supplement S1: Parameter and Variable Definitions

Supplement S2: Methods

Supplement S3: The HaloGen Simulation Framework

Supplement S4: Generalisation of the Effect of Increased Unknowns

In addition, details of all programs, user manuals and validation (continually updated) is available at https://sites.google.com/view/altrap/halogen and the Github site at https://github.com/peterdgill/HaloGen_Bayes

## 3. Results and Discussion

### 3.1. A Comprehensive and Flexible Framework for Activity-Level Evaluation

The HaloGen framework, detailed in this paper, represents a significant advance in the analysis of DNA quantities given activity-level propositions. It moves beyond the constraints of previous models, which were often limited to simpler scenarios, by creating a system that is both comprehensive and flexible. The core novelty of HaloGen lies in its ability to cohesively evaluate complex evidence sets involving multiple known or specified unknown contributors across any number of stains.

A key innovation is the treatment of background DNA. Rather than collapsing all non-specified individuals’ DNA into a single “background” parameter, HaloGen evaluates each specified unknown contributor as a distinct entity with its own associated DNA quantity evidence. This, combined with the use of a generalised fail-rate, *F*_0_, representing the probability that an unobserved actor leaves no detectable DNA, allows for an exhaustive evaluation. Furthermore, the framework’s modular design means that it is not static; it can be extended to incorporate new data and variables (Section 3.5). This creates a pathway towards a single, unified likelihood ratio (LR) that incorporates evidence from diverse forensic sources, such as shedder status or body fluid type, providing the court with a single, unified measure of evidential strength rather than a collection of disparate weights that are difficult to combine.

### 3.2. Interpreting LR Behaviour: Key Findings from Simulations

#### 3.2.1. Results of the Analysis

The theoretical principles of the framework are supported by results of simulated experiments, which demonstrate several key model behaviours. The data in Table 1 and Table 2 show the model’s sensitivity to the full context of the evidence, including DNA quantities, the number of stains, and the presence of other contributors. The framework models how the magnitude and variability of *q* affect the probability of the evidence under each proposition.

**Table 1:**
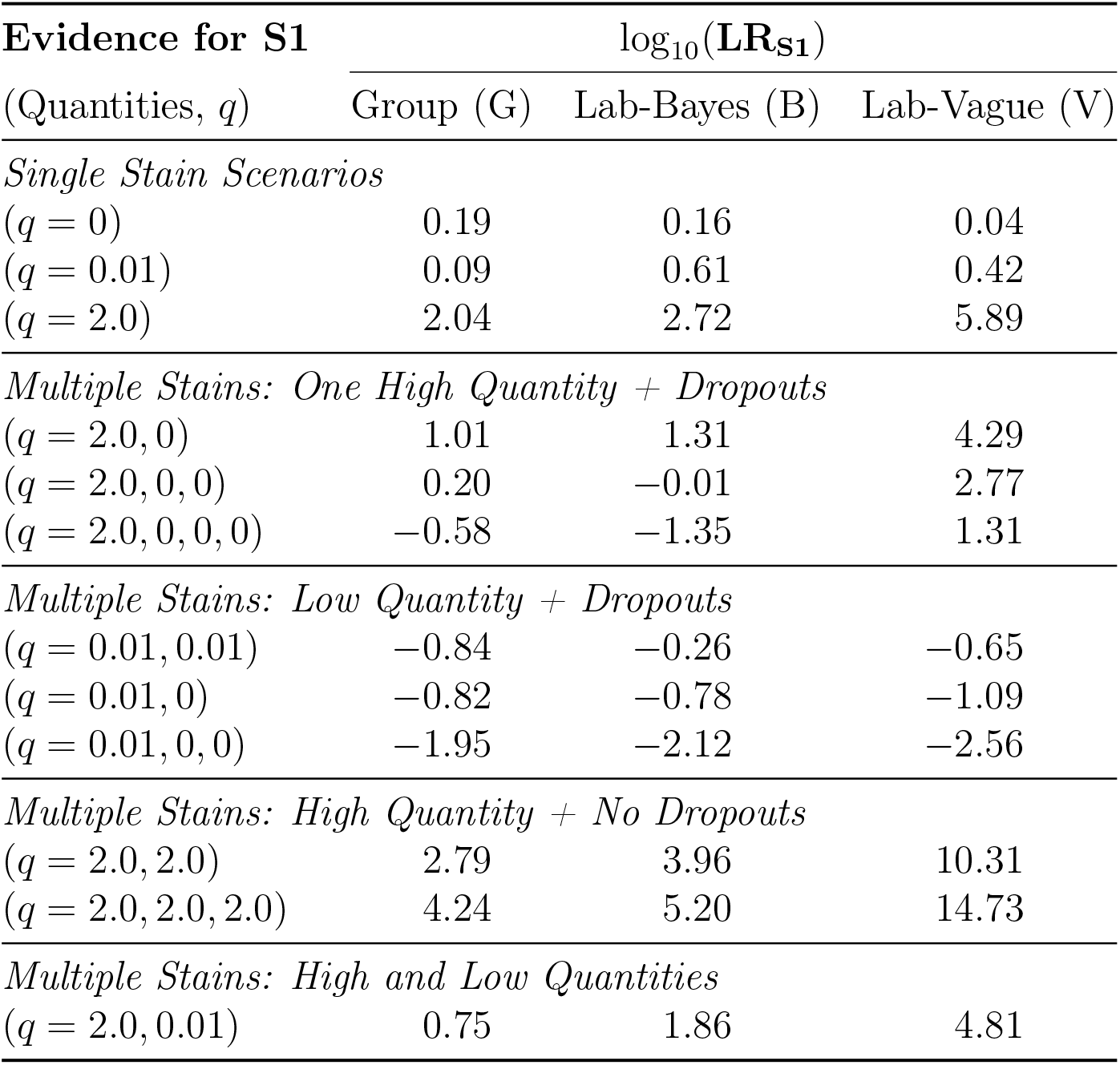
Likelihood Ratio (LR) results for S1 in single-contributor scenarios (*N*_*S*_ = 1, *n*_*U*_ = 0). LRs are presented as log_10_(LR) for the Group (G), Lab-Bayes (B), and Lab-Vague (V) models under the empirical-clamped *F*_0_ policy. Quantities are shown as vectors: e.g., *q* = 2.0, 0, 0 means three stains where 2.0 ng was recovered from the first stain and dropout was observed in the second and third stains. The value *q* = 0 denotes a non–detect, i.e. an observation at or below the detection limit DL. The calculation is not sensitive to the order.

**Table 2:** Likelihood Ratio (LR) results for scenarios with a known (S1) and a specified unknown (U1) contributor (*N*_*S*_ = 1, *n*_*K*_ = 1, *n*_*U*_ = 1). LRs are presented as log_10_(LR) for the Group (G), Lab-Bayes (B), and Lab-Vague (V) models. The top panel shows results for simple single-stain mixtures. The bottom panel shows results for more complex scenarios with two or three stains, where S1 and U1 may be present individually or in admixture.

| Evidence (Quantities) | | $\log_{10}(\mathbf{LR}_{\mathbf{S1}})$ | | | $\log_{10}(\mathbf{LR}_{\mathbf{U1}})$ | | |
| --- | --- | --- | --- | --- | --- | --- | --- |
| $q_{S1}$ | $q_{U1}$ | G | B | V | G | B | V |
| *Simple Scenarios (Single Stain Mixture)* |  |  |  |  |  |  |  |
| $(q = 0.01)$ | $(q = 0.01)$ | -0.26 | -0.10 | -0.14 | -0.26 | -0.10 | -0.14 |
| $(q = 0)$ | $(q = 0.01)$ | -0.10 | -0.64 | 0.61 | -0.38 | 0.24 | 0.09 |
| $(q = 0)$ | $(q = 2.0)$ | -1.78 | -2.75 | -6.09 | 1.57 | 2.38 | 5.56 |
| *Complex Scenarios (Multiple Stains)* |  |  |  |  |  |  |  |
| $(q = 0.01, 0)$ | $(q = 0.01, 0)$ | -0.88 | -0.85 | -1.12 | -0.88 | -0.85 | -1.12 |
| $(q = 0.01, 0)$ | $(q = 2.0, 0)$ | -2.77 | -2.23 | -5.52 | 0.93 | 1.19 | 4.22 |
| $(q = 2.0, 0, 0)$ | $(q = 0.01, 0, 0)$ | 0.10 | -0.02 | 2.76 | -3.17 | -2.78 | -5.71 |

##### Interpretation of *q* values in the tables

The outcomes shown in these tables represent hypothetical values of *q* under the generative model and include both detectable quantities (*q* ≥ DL) and non–detects (*q <* DL). In particular, the value *q* = 0 is used as a shorthand for a non–detect, meaning that either no DNA was transferred or that a positive amount fell below the detection limit.

#### 3.2.2. Single-Contributor Scenarios

Several consistent patterns emerge from Table 1.

1. **Baseline and low-quantity observations (***q* ≤ 0.01 **ng)**. With no or minimal DNA recovered, all models yield LR ≈ 1, typically favouring secondary transfer when low quantities appear repeatedly across stains. These results reflect appropriate down-weighting of near-threshold detections.
2. **Dropout sequences (e.g**. *q* = 2.0, 0, 0, …**ng)**. The LR decreases steadily as non-detections are added, indicating that repeated dropouts support the proposition of secondary transfer. The decline is proportional to the number of missing detections.
3. **Low-quantity stains with dropout (**0 *< q <* DL). All of these scenarios give low LR values, representing weak support for secondary transfer.
4. **High-quantity observations (***q* = 2.0 **ng)**. A single strong stain provides support for direct transfer. When several high-quantity stains are observed, progressively higher LRs are observed.
5. **Mixed patterns (high and low quantities)**. When both strong and weak stains occur together, the overall weight of evidence is dependent upon relative contributions of DNA quantities.

##### Summary and dependency note

HaloGen responds predictably to both the magnitude of recovered DNA and the pattern of detections across stains. High quantities consistently support direct transfer, while weak or inconsistent evidence lead to neutral or outcomes supporting secondary transfer. This stability results from the empirical-clamped *F*_0_ policy, which bounds probabilities within data-informed limits [*J*, 1 − *J*] and prevents artificial inflation of likelihood ratios in low-information cases.

Importantly, all stains for a contributor are analysed jointly within a single Bayesian model. HaloGen constructs a *joint likelihood* across all stains from that contributor, linking them through shared latent parameters (*µ, σ, k*). The stains are therefore *conditionally independent* given these parameters, but become statistically dependent once uncertainty in (*µ, σ, k*) is considered. Each additional stain contributes to updating the joint likelihood for a given contributor, and the resulting likelihood ratio reflects the weight of the combined evidence, instead of a simple product of independent terms.

#### 3.2.3. Analysis of Multi-Contributor Scenarios (Table 2)

The scenarios involving both S1 and U1 illustrate three core behaviours of the framework: (i) the *contributor-specific* nature of anchoring, (ii) coherent reallocation of weight when an alternative specified contributor shows a detected quantity, and (iii) explicit control of extreme values via the empiricalclamped *F*_0_ policy.

##### Contributor-specific anchoring (simple, low-level quantities)

When both S1 and U1 show only small quant values on single stains (*q*_*S*1_ = 0.01, *q*_*U*1_ = 0.01), all models return weak support for secondary transfer for each contributor considered in isolation (S1: − 0.26, − 0.10, − 0.14; U1: − 0.26, − 0.10, − 0.14). This reflects that small detected quantities are, *a priori*, more typical of secondary mechanisms than of direct transfer for the fitted posteriors.

When S1 is a dropout but U1 has a small quant (*q*_*S*1_ = 0, *q*_*U*1_ = 0.01), the LR for *U1* is near-neutral to slightly supportive of direct transfer, depending on the model (Group: −0.38; Lab-Bayes: +0.24; Lab-Vague: +0.09), whereas the LR for S1 weakly supports proposition of secondary transfer across models (S1: −0.10, −0.64, −0.61). This demonstrates contributor-specific anchoring behaviour: the detected quantity anchors inference for U1, while S1 (having only a dropout) does not benefit from an evidential anchor and therefore cannot support the proposition of direct transfer.

##### Reallocation of weight under a strong alternative (simple, high quantity)

If S1 is a dropout and U1 shows a high quantity (*q*_*S*1_ = 0, *q*_*U*1_ = 2), the framework assigns strong support for direct transfer to U1 and strong support for secondary transfer to S1. Numerically, S1 log_10_ LR is negative across all models (− 1.78, − 2.75, − 6.09), while U1 supports proposition of direct transfer (+1.57, +2.38, +5.56). This behaviour is expected: a high observed quantity is far more probable under direct transfer for U1 (the alternative contributor) than under S1’s direct-transfer pathway when S1 exhibits no detection, so the denominator of LR_S1_ dominates.

##### Complex evidence patterns (multiple stains)

With two stains showing low-level signals for both contributors (*q*_*S*1_ = 0.01, 0.00; *q*_*U*1_ = 0.01, 0.00), the results remain mildly defence-leaning and symmetric for each contributor (S1: −0.88, −0.85, −1.12; U1: −0.88, −0.85, −1.12), consistent with weak, threshold-adjacent detections plus dropouts.

When S1 shows a small quant plus a dropout while U1 shows a high quant plus a dropout (*q*_*S*1_ = 0.01, 0.00; *q*_*U*1_ = 2, 0), S1’s log_10_ LR is clearly negative (−2.77, −2.23, −5.52), whereas U1’s is clearly positive (+0.93, +1.19, +4.22). A detected high quantity for U1 provides a strong anchor for that contributor and simultaneously explains why S1’s LR falls below unity under the competing denominator hypotheses.

Finally, when S1 shows a high quantity together with multiple non-detects and U1 shows only a small quant with dropouts (*q*_*S*1_ = 2, 0, 0; *q*_*U*1_ = 0.01, 0.00, 0.00), the overall LR result for S1 reduces as the number of dropouts increase (S1: +0.10, −0.02, +2.76), while U1 remains strongly disfavoured (U1: −3.17, −2.78, −5.71).

When a high quantity of DNA is obtained from a contributor, the LR value is moderated by background or secondary DNA transfer and repeated non-detections,

#### 3.2.4. Generalising the Impact of F_0_: A Sensitivity Analysis

To illustrate the effect of the fail-rate parameter in the most transparent setting, we return to the simplest case considered earlier: a single contributor and a single actor (section 2.5.1).

It is informative to visualise how the likelihood ratio (LR) behaves as a function of *F*_0_ in the simplified one-contributor case to understand its impact. For a fixed evidence-strength ratio *C* = *t*_1_*/s*_1_, the theoretical relationship is

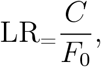

showing that *F*_0_ for an unobserved actor directly scales the final LR. Here, *C* represents the intrinsic strength of the DNA evidence: how many times more probable the observed quantity is under direct transfer (*H*_*p*_) than under secondary or background transfer (*H*_*d*_). In the full HaloGen inference, *C* and *F*_0_ are jointly determined from the same posterior draws of the experimental model; here, *C* is held fixed purely to isolate and visualise the marginal effect of *F*_0_.

Figure 2 illustrates this sensitivity, plotting LR versus *F*_0_ for several fixed *C* values. The curves represent theoretical envelopes for how the LR would change across all *F*_0_ values.

**Figure 2:**
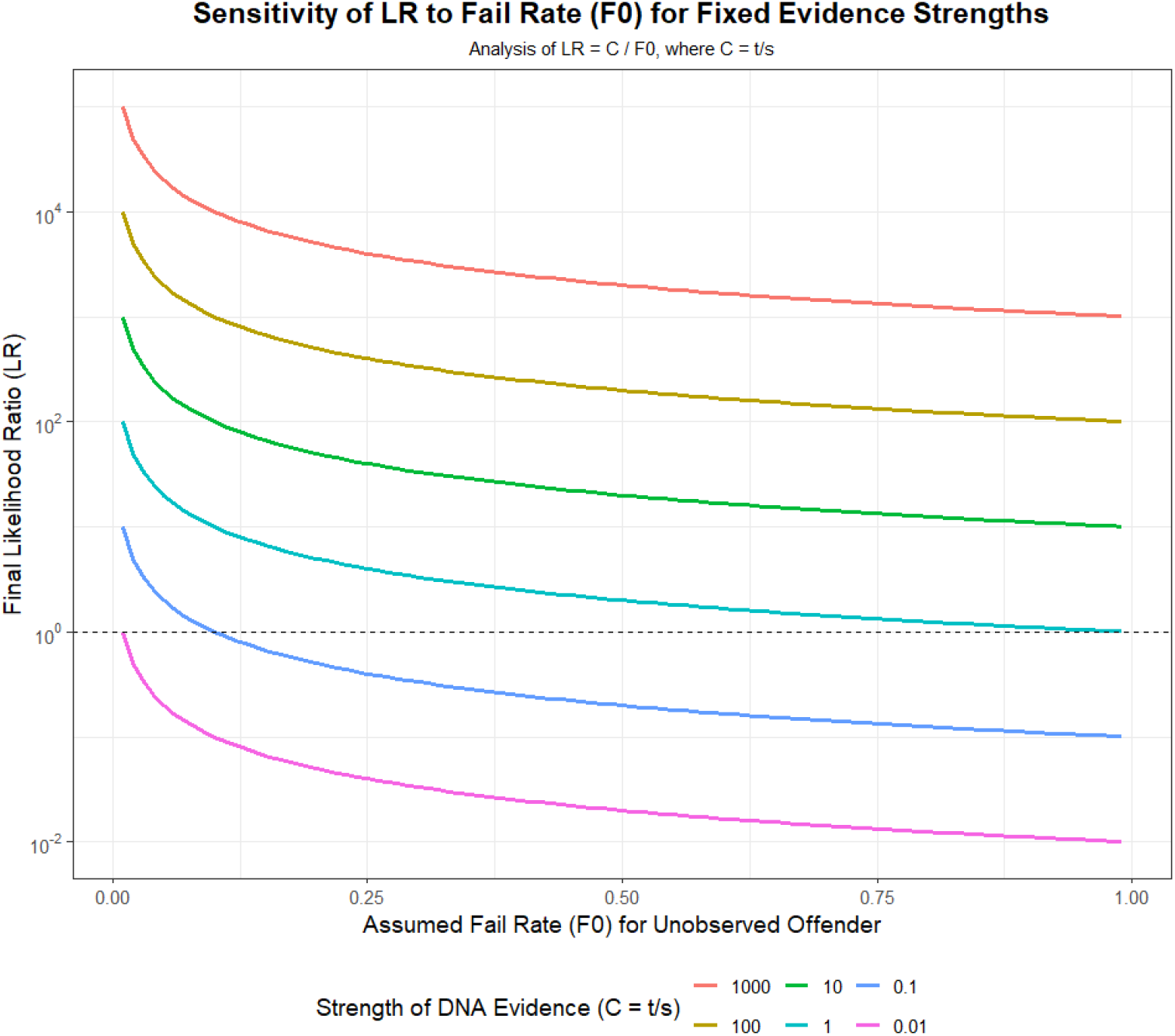
Sensitivity of the final LR to the assumed fail rate (*F*_0_) for different fixed evidence strengths (*C* = *t/s*). The LR axis is shown on a logarithmic scale.

1. **Inverse proportionality**. The LR decreases inversely with *F*_0_. For any fixed evidence strength *C*, increasing the probability that an unobserved actor leaves no detectable trace reduces the evidential weight against the specified contributor, often by several orders of magnitude as *F*_0_ approaches one.
2. **Dominance of the fail rate (***F*_0_). The fail rate (*F*_0_) can dominate the overall LR. Even when the DNA evidence itself is strong (e.g. *C* = 1000, meaning *t*_1_ is 1000 times greater than *s*_1_), a plausible *F*_0_ of 0.1 reduces log_10_(LR) by roughly one unit compared with *F*_0_ = 0.01. This underscores why HaloGen bounds *F*_0_ empirically per draw: preventing unrealistically small values that would otherwise inflate LRs.
3. **Contextual importance**. Reporting an LR based solely on *t* and *s* is incomplete without explicitly addressing *F*_0_. The conclusion can be highly sensitive to the assumed probability that an unobserved actor leaves no trace.
4. **Limiting behaviour when** *t* = *s*. For completeness, note that when *C* = 1 (i.e. the observed quantity is equally probable under direct and secondary transfer), the LR reduces to LR = 1*/F*_0_. In this limiting case LR = 1 only when *F*_0_ = 1, which corresponds to a degenerate scenario in which an actor always leaves no detectable DNA. This boundary case serves only as a reference and does not represent a realistic forensic situation.

Future refinements may allow *F*_0_ to vary with contextual factors such as surface type, substrate, or time since deposition.

This sensitivity analysis provides the theoretical basis for the *empirical–clamped policy* described in Section 2.3.2, ensuring that the *F*_0_ term cannot take unrealistically small values that would spuriously inflate support for direct transfer.

The current implementation uses a common empirically constrained *F*_0_ for the relevant unobserved actor within a given model and laboratory. This avoids introducing an unvalidated asymmetry between propositions. In principle, however, *F*_0_ could be proposition-specific if the competing propositions imply materially different opportunities for deposition, persistence, recovery or sampling. Such an extension would require explicit empirical support or a clearly disclosed proxy, because proposition-specific *F*_0_ values can have a substantial effect on the LR. Where such differences are plausible but not well quantified, sensitivity analysis should be reported rather than a single unqualified LR.

#### 3.3. The ‘Anchor Principle’: A Guideline for Meaningful Application

The results in Tables 1 and 2 show a key interpretive safeguard in HaloGen, referred to as the *anchor principle*. Simulations show that a single stain with no DNA recovered (*q* = 0) can sometimes yield a very weak apparent value (LR_POI_ *>* 1) supporting direct transfer (Table 2). This outcome is not paradoxical. It arises because the model compares two small probabilities: the probability of observing no DNA under *H*_*p*_: (*k*_*D*_ +(1 − *k*_*D*_)*F*_*D*_(DL)) versus under *H*_*d*_: (*F*_0_ ≈ the same value). When the estimated probability of dropout under the direct model is similar in magnitude to the assumed fail rate *F*_0_, the ratio can be slightly greater than one. From an interpretive standpoint, however, such results are not meaningful. An empty stain provides no genetic evidence that can be attributed to a specific individual. The empirical-clamped policy ensures that such cases remain neutral in practice.

The multi-contributor simulations reinforce this principle (Table 2). When one contributor (e.g. S1) has a dropout (*q*_*S*1_ = 0) while another (U1) yields a small detectable quantity (*q*_*U*1_ = 0.01), the model returns a meaningful LR for U1 but a negative log_10_(LR) for S1. This is the expected behaviour: the detected quantity provides an *anchor* for U1’s inference, whereas S1’s absence of signal carries no probative weight.

This leads to a practical interpretive rule: *A quantitative likelihood ratio for a specific person of interest (POI) is only meaningful when the evidence set provides an anchor; that is, at least one detected, non-zero DNA quantity attributable to that person*. Without such a contributor-specific anchor, the quantitative evidence for that person should be regarded as uninformative and reported as neutral (LR ≈ 1). This ensures that the weight of evidence reflects what has actually been observed, not hypothetical absences.

### 3.4. Implications for Casework and the Avoidance of Confirmation Bias

The HaloGen framework has significant implications for forensic casework. By construction, it requires the analyst to evaluate *all* specified contributors (both known and unknown) across *all* relevant stains (positive and negative results) simultaneously. This exhaustive approach acts as a procedural safeguard against *confirmation bias*, the tendency to focus selectively on findings that support a favoured hypothesis or person of interest (POI) while discounting conflicting information. Because every contributor is explicitly represented in every elemental hypothesis, and a likelihood ratio (LR) can be computed for each, HaloGen enables balance and transparency in the interpretation process. As illustrated by the Tengs case [19], overlooking DNA signals from unknown contributors can profoundly distort the inferential picture; HaloGen’s exhaustive design mitigates this risk.

The framework also invites reflection on casework sampling strategies. Decisions about where and how to sample—for example, targeted swabbing of likely contact areas, contextual/background sampling, or more extensive sampling of associated surfaces—produce data with different inferential implications. Recent work on background and contextual sampling has emphasised that the number, size, and location of samples affect the probabilities assigned at activity level and can provide case-relevant information about prevalence and background [20, 21]. Within HaloGen, non-detections can be incorporated as part of the evidence set when they arise from a sampling strategy that is relevant to the propositions. However, determining what constitutes a relevant area to test, and interpreting non-detections when contact itself is uncertain, remain case-specific scientific judgements rather than automatic outputs of the software.

The Tengs case provides a practical illustration. Approximately sixty swabs were taken from a single item of clothing (the pantyhose). Two yielded results aligning with the defendant; roughly four indicated unknown contributors; and the remainder produced no interpretable DNA. The evidential question is how to reconcile this heterogeneous mixture of detections and dropouts into a coherent probabilistic statement. A full application of the HaloGen framework to such a dataset lies beyond the scope of this theoretical paper, but the case underscores a key point: a comprehensive, model-based approach enables the joint evaluation of all samples and outcomes, providing a principled alternative to *ad hoc* or selective interpretation.

### 3.5. Future Extensions: Incorporating Shedder Variability

A major strength of the HaloGen framework is its modular architecture, which allows new sources of biological variability to be incorporated without altering the core structure. One important future extension, planned for development once suitable experimental data are available, is the explicit modelling of *individual shedder variability*.

It is well established that individuals differ in the amount of DNA they deposit during contact. Within the HaloGen framework, this variability can be represented as a person-specific random effect that modulates the expected DNA quantity for any given transfer process. Formally, the mean log-quantity parameter for a stain, *µ*, can be decomposed as

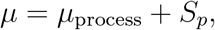

where *µ*_process_ captures the baseline effect of the transfer activity (e.g., direct or secondary), and *S*_*p*_ represents the individual’s relative shedding propensity.

The shedder effect *S*_*p*_ would be treated as a latent random variable drawn from a population-level distribution centred at zero,

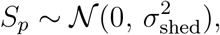

so that *σ*_shed_ quantifies the degree of inter-individual variability in DNA deposition rates. This parameter could be estimated empirically from controlled shedding experiments involving multiple participants. For individuals studied repeatedly, the same structure could also capture within-person variation across time or activity type.

Once characterised, the shedder term can be integrated directly into HaloGen’s hierarchical Bayesian model. In calculating a likelihood ratio, the framework would then marginalise not only over the posterior uncertainty in transfer parameters (*µ*_process_, *σ, k*) but also over the unobserved shedder effects *S*_*p*_ for the contributors involved. This extension naturally embeds shedder variability into the model’s generative process, rather than treating it as an external adjustment or qualitative consideration.

Beyond shedding, the same principle provides a blueprint for incorporating other biologically relevant factors, such as substrate type, body fluid origin, or time since deposition, within a unified probabilistic structure. The overarching goal is to deliver to the court a single, transparent likelihood ratio that integrates multiple interacting sources of uncertainty, rather than a collection of separate, disconnected statistics. This model-first approach not only enhances interpretability but also identifies which experimental parameters are most informative, thereby guiding the design of future empirical studies.

### 3.6. Model Assumptions and Limitations

While the HaloGen framework provides a comprehensive and flexible quantitative approach, its application requires awareness of its underlying assumptions Like all probabilistic models, it offers a structured representation of complex biological and procedural processes.

1. **Dependence on Data Quality and Representativeness:** The reliability of the likelihood ratios (LRs) produced by HaloGen depends on the quality, quantity, and relevance of the experimental data used to estimate its core parameters (*µ, σ, k*). If the calibration data do not accurately reflect the materials, substrates, transfer mechanisms, or laboratory protocols relevant to a given case, the resulting LRs may be unrepresentative. High measurement uncertainty, limited sample sizes, or systematic bias in the reference data can propagate directly into the case-level inference.
2. **Primacy of Hypothesis Formulation:** HaloGen computes *P* (*E* | *H*) for any well-defined proposition *H*, but it does not determine what those propositions should be. The forensic expert must articulate case-appropriate hypotheses (*H*_*p*_ and *H*_*d*_), specify the conditioning (e.g., number of actors, *N*_*S*_), and define the relevant activities. The validity of any LR ultimately depends on the validity of the propositions it compares; the framework cannot correct for ill-posed or contextually inappropriate hypotheses.
3. **Core Modelling Assumptions:** The model assumes that observed DNA quantities follow a a *zero-inflated and censored lognormal distribution*, which explicitly accounts for the analytical detection limit (DL) and the possibility of true dropouts. This form has been found empirically to fit well.

Alternative forms could be adopted if warranted by future data.

The fail-rate parameter *F*_0_, representing the probability that an unobserved actor leaves no *detectable* trace, in the current implementation is informed by the empirical distribution of direct-transfer dropout rates. This pragmatic assumption reflects the idea that unobserved actors behave similarly to measured direct transfers but can be revisited as more empirical evidence accumulates. Sensitivity analysis is important. Section 3.2.4 examines how varying this assumption affects the resulting LRs. In future work, *F*_0_ may be allowed to vary across contributors or case types.

1. **Scope limitation: the anchor principle**. As discussed in Section 3.3, the model is not intended for application to evidence sets that produce no detectable amount of contributor-level DNA. A meaningful contributor-specific activity-level LR requires an *evidential anchor* : at least one observed, quantifiable DNA result that can be attributed to the contributor with sufficient sub-source support. In the absence of such an anchor, HaloGen should not be used to generate a positive activity-level LR for that person.

For reporting, the following rules are recommended. First, a contributor-specific activity-level LR should be reported only where there is at least one detected, quantifiable contributor-level DNA result that can be attributed to that contributor with sufficient sub-source support. Second, if a person has no detectable contributor-level quantity, the appropriate conclusion is neutral or not evaluable given activity level, unless the propositions specifically concern the absence of DNA and the relevant non-detect probabilities are supported by suitable data. Third, if contributor-level quantities cannot be reliably assigned across stains, this limitation should be stated explicitly, and the analysis should either incorporate that uncertainty upstream or be restricted to propositions that the data can support.

In summary, the HaloGen framework offers transparency and internal consistency but does not remove the need for critical judgment. Its validity depends on the inputs and the appropriateness of its use within the defined evidential context.

## 4. Conclusion

This paper has detailed the complete theoretical basis for the HaloGen framework, a next-generation system for the evaluation of DNA quantity evidence in complex, activity-level scenarios. We have presented the mathematical formulations for handling multiple contributors and stains. The concept of the number of actors is crucial since this links to the role of the generalised fail rate *F*_0_, and the logic behind interpreting the model’s output. This culminates in the formalisation of the ‘anchor principle’. The theory demonstrates that HaloGen is a flexible, transparent, and logically robust framework that avoids common sources of bias by integrating all available information. The companion paper, Part 2 [18], will explore the practical characterisation and validation of the model, providing recommendations for its adoption in casework.

## Acknowledgements

Funding for this Project was received from the European Union’s Internal Security Fund -Police (ISFP) – with Grant Agreement title: Competency, Education, Research, Testing, Accreditation, and Innovation in Forensic Science” [CERTAIN-FORS] ISFP-2020-AG-IBA-ENFSI and number: 101051099. Also: ReAct II – Extension of ReAct. Work package 5 within the FOR-FUTURE project - Forensic Fundamentals, Technology, Multidisciplinarity, Research, Evaluation (FOR-FUTURE) — ISF-2023-TF2-AG-ENFSI-IBA-2

## References

[1] P. Gill, T. Hicks, J. M. Butler, E. Connolly, L. Gusmão, B. Kokshoorn, N. Morling, R. A. van Oorschot, W. Parson, M. Prinz, et al., DNA commission of the International society for forensic genetics: Assessing the value of forensic biological evidence-Guidelines highlighting the importance of propositions. Part II: Evaluation of biological traces considering activity level propositions, Forensic Science International: Genetics 44 (2020) 102186.

[2] P. Gill, A. E. Fonneløp, T. Hicks, S. Xenophontos, M. Cariolou, R. van Oorschot, I. Buckel, V. Sukser, S. Papić, S. Merkaš, et al., The ReAct project: Analysis of data from 23 different laboratories to characterise DNA recovery given two sets of activity level propositions, Forensic Science International: Genetics 76 (2025) 103222.

[3] P. Gill, T. Hicks, A. Carracedo, The ReAct project: Bayesian networks for assessing the value of the results given activity level propositions, Forensic Science International: Genetics 76 (2025).

[4] C. G. G. Aitken, F. Taroni, S. Bozza, The Role of the Bayes Factor in the Evaluation of Evidence, Annual Review of Statistics and Its Application 11 (1) (2024) 203–230.

[5] A. Gelman, Multilevel (hierarchical) modeling: what it can and cannot do, Technometrics 48 (3) (2006) 432–435.

[6] C. G. G. Aitken, Bayesian Hierarchical Random-Effects Models in Forensic Science, Frontiers in Genetics 9 (2018) 126.

[7] D. Taylor, L. Volgin, P. Gill, B. Kokshoorn, Accounting for interlaboratory DNA recovery data variability when performing evaluations given activities, Forensic Science International: Genetics 78 (2025) 103283.

[8] Ø. Bleka, G. Storvik, P. Gill, EuroForMix: An open-source software based on a continuous model to evaluate STR DNA profiles from a single source and mixtures, Forensic Science International: Genetics 21 (2016) 35–44.

[9] J.-A. Bright, D. Taylor, J. M. Curran, J. S. Buckleton, Developing allelic and stutter peak height models for a continuous method of DNA interpretation, Forensic Science International: Genetics 7 (2) (2013) 296–304.

[10] H. Haned, P. Gill, C. Poot, T. Egeland, P. Stoop, Development and validation of a statistical model to evaluate the probative value of quantitative DNA results in ‘he-said-she-said’ cases, Forensic Science International: Genetics 22 (2016) 207–216.

[11] L. Samie, R. van Oorschot, DNA transfer: A review, Wiley Interdisciplinary Reviews: Forensic Science 4 (1) (2022) e1435.

[12] F. Taroni, S. Bozza, A. Biedermann, P. Garbolino, C. Aitken, Data Analysis in Forensic Science: A Bayesian Decision Perspective, John Wiley & Sons, Chichester, UK, 2010.

[13] T. Hicks, Z. Kerr, S. Pugh, J.-A. Bright, J. Curran, D. Taylor, J. Buckleton, Comparing multiple POI to DNA mixtures, Forensic Science International: Genetics 52 (2021) 102481.

[14] K. Slooten, The comparison of DNA mixture profiles with multiple persons of interest, Forensic Science International: Genetics 56 (2022) 102592.

[15] B. Carpenter, A. Gelman, M. D. Hoffman, D. Lee, B. Goodrich, M. Betancourt, M. Brubaker, J. Guo, P. Li, A. Riddell, Stan: A Probabilistic Programming Language, Journal of Statistical Software 76 (1) (2017) 1–32.

[16] Ø. Bleka, M. D. Vigeland, P. Gill, EFMex: Using EuroForMix to evaluate DNA mixtures with multiple persons of interest, Forensic Science International: Genetics (2025) 103365.

[17] P. Gill, T. Hicks, B. Kokshoorn, R. A. van Oorschot, D. Taylor, W. Parson, Minimum FSI: Genetics requirements for publishing data on DNA transfer and recovery, given activities, Forensic Science International: Genetics (2025) 103330.

[18] P. Gill, Ø. Bleka, Likelihood Ratios Given Activity-Level Propositions for DNA Transfer Evidence: Practical Implementation and Simulation Studies Using the HaloGen Engine (Part II), bioRxiv (2026). doi: 10.64898/2026.02.06.703509. URL https://www.biorxiv.org/content/10.64898/2026.02.06.703509v1

[19] P. Gill, M. M. Andersen, J. P. Whitaker, A. J. Kal, W. Parson, Birgitte Tengs case: analysis and the wider implications for evaluation of DNA evidence given activities, Forensic Science International: Genetics (2025) 103279.

[20] J. B. Reither, D. Taylor, B. Szkuta, R. A. van Oorschot, Determining the number and size of background samples derived from an area adjacent to the target sample that provide the greatest support for a POI in a target sample, Forensic Science International: Genetics 68 (2024) 102977.

[21] Y. R. Goedhart, J. A. de Koeijer, L. H. Aarts, C. J. de Poot, B. Kokshoorn, Contextual DNA samples in evaluations given activity level propositions, Forensic Science International: Genetics 84 (2026) 103452.

