## Supplementary Files for "Likelihood Ratios Given Activity-Level Propositions for DNA Transfer Evidence: Theoretical Foundations of the HaloGen Framework (Part I)"

#### S1. Parameter and Variable Definitions

This glossary standardises the notation used across the HaloGen framework.

*I. Scenario and Inputs*

$E_c$ Let $E_c$ denote the detected, quantifiable contributor-level evidence for contributor $c$, unless a section explicitly states that non-detect observations are being modelled. Thus, in the main case-level LR construction, $E_c$ consists of observed quantities $q_{c,j} \geq$ DL that can be attributed to contributor $c$ with sufficient sub-source support. Non-detect observations are treated separately through probability-mass terms, including $F_0$ where an elemental hypothesis requires an unobserved actor.

$q_c$, $q_i$ Observed DNA quantity attributed to contributor $c$; $q_i$ denotes stain $i$.

$DL$ Detection limit (below this, values are treated as non-detects).

$n_K$ Number of specified known contributors.

$n_U$ Number of specified unknown contributors.

$n$ Total specified contributors, $n = n_K + n_U$.

$N_S$ Assumed number of relevant actors in the case.

*II. Per-Transfer Statistical Model (DL-aware)*

Each transfer type (direct $D$ or secondary $S$) is modelled by a zero-inflated, left-censored Lognormal mixture.

$\mu$ Mean of $\log(q)$ for the Lognormal component.

$\sigma$ Standard deviation of $\log(q)$.

$k$ 'no-transfer event' probability: Pr(no detectable DNA in experiment).

$(\mu_D, \sigma_D, k_D)$ Direct-transfer parameters.

$(\mu_S, \sigma_S, k_S)$ Secondary-transfer parameters.

$f_{\mathrm{LN}}(q; \mu, \sigma)$ Lognormal density for $q > 0$.

$F(\cdot)$ Lognormal cumulative distribution function.

$z_0$ Standardised detection-limit point: $z_0 = (\log DL - \mu)/\sigma$.

*DL-aware experimental likelihood.*

$$L(q \leq DL \mid \theta, DL) = k + (1-k)F(DL), \qquad L(q > DL \mid \theta, DL) = (1-k) f_{\mathrm{LN}}(q; \mu, \sigma).$$

*III. Hierarchical (Group) Hyperparameters*

$\mu_0,\ \tau_\mu$ Population mean and SD of lab-level $\mu$.

$\log \sigma_0,\ \tau_{\log \sigma}$ Population mean and SD of lab-level $\log \sigma$.

$\mu_k,\ \phi_k$ Mean and precision of the population Beta prior for $k$.

*IV. Lab-Specific Parameters*

$\mu_{\mathrm{lab}}$, $\log \sigma_{\mathrm{lab}}$, $k_{\mathrm{lab}}$ Lab-specific parameters under Lab–Bayes or Lab–Vague.

$\theta$ Generic parameter vector $(\mu, \sigma, k)$ for a given transfer type.

*V. Contributor-Wise Posterior–Predictive Likelihoods*

For any contributor $c$ with detected stains $\mathbf{q}_c$:

$t_c$ Posterior–predictive likelihood under the *direct* transfer model.

$s_c$ Posterior–predictive likelihood under the *secondary/background* model.

$\tau'_j, s'_j$ Direct and secondary likelihoods for specified unknown contributor $u_j$.

*Case-level detect-side rule (unified conditional form).* Consistent with Section 2.3:

- For any *detected* stain ($q > DL$), both $t_c$ and $s_c$ use the truncated density

$$L(q > DL \mid \theta, DL) = \frac{f_{\mathrm{LN}}(q; \mu, \sigma)}{1 - F(DL)}.$$

- For *non-detects* ($q \le DL$):

$$L(q \le DL \mid \theta, DL) = k + (1 - k)F(DL).$$

Thus both transfer types use the same conditional-on-detection form at case level (as described in Section 2.3).

*VI. Exhaustive Elemental Propositions*

$\mathcal{K}, \mathcal{U}$ Sets of known individuals and specified unknown contributors.

$\mathcal{O} \subseteq \mathcal{K} \cup \mathcal{U}$ Subset of specified contributors assigned to the direct-transfer actor role under elemental hypothesis $H_{\mathcal{O}}$.

$x$ Number of specified contributors assigned to the direct-transfer actor role:

$$x = |\mathcal{O}|.$$

*Elemental likelihood (Section 2.4).* For an elemental hypothesis $H_{\mathcal{O}}$, the contributors in $\mathcal{O}$ are evaluated using direct-transfer likelihood components, whereas contributors in $(\mathcal{K} \cup \mathcal{U}) \setminus \mathcal{O}$ are evaluated using secondary-transfer likelihood components. If the activity-level proposition requires $N_S$ actors in total, then $N_S - x$ actors are unobserved and contribute through the case-level non-detection probability $F_{0,t}$:

$$P(E \mid H_{\mathcal{O}}, N_S) = \left( \prod_{c \in \mathcal{O}} t_c \right) \left( \prod_{c \in (\mathcal{K} \cup \mathcal{U}) \setminus \mathcal{O}} s_c \right) (F_{0,t})^{(N_S - x)}.$$

Equivalently, since $x = |\mathcal{O}|$,

$$P(E \mid H_{\mathcal{O}}, N_S) = \left( \prod_{c \in \mathcal{O}} t_c \right) \left( \prod_{c \in (\mathcal{K} \cup \mathcal{U}) \setminus \mathcal{O}} s_c \right) (F_{0,t})^{(N_S - |\mathcal{O}|)}.$$

Only admissible elemental hypotheses with $|\mathcal{O}| \le N_S$ are included.

*VII. Unobserved Actor Term*

$F_0$ Case-level probability that a relevant but unobserved actor leaves no detectable DNA.

**Empirical–Jeffreys update** $F_0 \sim \text{Beta}(f + 0.5, n - f + 0.5)$ with mean $J = (f + 0.5)/(n + 1)$.

**Clamp** Each draw is constrained to $[J, 1 - J]$.

**Tail component** $F_{0,t}^{\text{tail}} = k_D^{(t)} + (1 - k_D^{(t)})F_D(\text{DL})$.

**Effective case-level non-detect probability** $F_{0,t} = \max(F_{0,t}^{\text{policy}}, F_{0,t}^{\text{tail}})$.

*VIII. Miscellaneous*

$m$ Number of replicate stains for contributor $c$.

LR Likelihood ratio. For each posterior Monte Carlo draw, HaloGen computes an LR/BF draw from the paired direct-transfer and secondary-transfer likelihood components. The reported point estimate is the median of these LR/BF draws, with uncertainty summarised by the 10–90% quantiles.

$S_p, \sigma_{\textbf{shedding}}$ Optional shedding random effect and its population SD.

### S2. Methods Supplement

This supplement provides technical details that underpin the conceptual description in the main Methods Section 2. A complete glossary of all parameters, variables, and notation is given in S1.

#### *S2.1. Case-Level $F_0$: Policy, Tail Derivation, and Use*

In the main text, Section 2.3.1 introduces the case-level parameter $F_0$ as the probability that an actor leaves no *detectable* DNA. This section details the operational policy used in HaloGen.

*Direct tail probability from the experimental model.* For direct transfer, the DL-aware experimental model in Section 2.1 yields parameters $\theta_D^{(t)} = (\mu_D^{(t)}, \sigma_D^{(t)}, k_D^{(t)})$ for each posterior draw $t$. The corresponding direct-transfer non-detect probability is

$$F_{0,t}^{\text{tail}} = k_D^{(t)} + \big(1 - k_D^{(t)}\big) F_D\big(DL \mid \mu_D^{(t)}, \sigma_D^{(t)}\big),$$

where $F_D(\cdot)$ is the Lognormal CDF (see Section 2.1). This quantity is the experimental analogue of "actor contribution not detected" for direct transfer.

*Jeffreys-smoothed empirical policy.* Let $x$ denote the number of non-detects and $n$ the total number of direct-transfer experiments. HaloGen defines a Beta distribution

$$F_0^{\text{policy}} \sim \text{Beta}(x + 0.5,\ n - x + 0.5)$$

with mean

$$J = \frac{x + 0.5}{n + 1}.$$

Each Monte Carlo draw from this Beta distribution is clamped to the interval $[J,\ 1 - J]$,

$$F_{0,t}^{\text{policy}} \leftarrow \min\Big\{1 - J,\ \max\{J,\ F_{0,t}^{\text{policy}}\}\Big\},$$

to avoid extreme values (0 or 1) that would otherwise produce numerically unstable or overly decisive likelihood ratios.

*Extra-cautious combination.* To remain defence-conservative, HaloGen combines the direct tail and the empirical policy via

$$F_{0,t} = \max\big\{F_{0,t}^{\text{policy}},\ F_{0,t}^{\text{tail}}\big\}.$$

This ensures that the case-level non-detect probability is never smaller than either the experimental-based tail or the empirical Beta estimate.

*Use in elemental hypotheses.* In the exhaustive LR construction (Section 2.4), if an elemental hypothesis $H_{\mathcal{O}}$ requires $N_S - |\mathcal{O}|$ unobserved actors, the contribution from these undetected actors is

$$(F_{0,t})^{N_S - |\mathcal{O}|}.$$

This is the same quantity labelled $F_0$ in the main text of Section 2.4; here we show its draw-wise construction explicitly.

*S2.2 Relationship Between Experimental $k$ and Case-Level $F_0$*

Section 2.1 defines the DL-aware experimental model, with parameters $\theta = (\mu, \sigma, k)$. For each transfer type, the experimental non-detect probability at the laboratory detection limit $DL$ is

$$P(q < DL \mid \theta) = k + (1 - k)\, F(DL),$$

where $F(DL)$ is the Lognormal CDF (Section 2.1).

Conceptually, $k$ and $F_0$ play different roles:

- $k$ is an *experimental-level* parameter: the probability that a single experimental transfer event yields no measurable DNA under controlled lab conditions ('no-transfer event').
- $F_0$ is a *case-level* quantity (Section 2.3.1): the probability that an *unobserved actor* in the case leaves no detectable DNA, given the broader case circumstances.

In practice, the estimation of $F_0$ is informed by the same experimental process, the direct-transfer DL-aware model, through the tail-based non-detect probability $F_{0,t}^{\text{tail}} = k_D^{(t)} + (1 - k_D^{(t)}) F_D(DL)$, subsequently combined with the empirical Beta policy. Thus, $F_0$ and $k$ are numerically linked but conceptually distinct: $k$ remains purely experimental ('no-transfer event'), whereas $F_0$ is the case-level parameter of non-detection that used in the exhaustive LR formulation.

*S2.3. Statistical Specification of Group, Lab-Bayes, and Lab-Vague Models*

Section 2.2 in the main text describes the three model families (*Group*, *Lab–Bayes*, *Lab–Vague*) conceptually. Here we provide the corresponding statistical specification.

*S2.3.1 Group model (hierarchical).* For each laboratory $l$, we consider summary parameters $(\hat{\mu}_l, \widehat{\log \sigma_l}, \hat{k}_l)$ derived from DL-aware fits to that lab's raw data (Section 2.1). The Group model assumes:

$$\hat{\mu}_l \sim \mathcal{N}(\mu_0, \tau_\mu^2), \qquad \widehat{\log \sigma_l} \sim \mathcal{N}(\log \sigma_0, \tau_{\log \sigma}^2),$$
$$\hat{k}_l \sim \text{Beta}\big(\mu_k \phi_k,\ (1 - \mu_k)\phi_k\big),$$

with weakly informative hyperpriors,

$$\mu_0, \log \sigma_0 \sim \mathcal{N}(0, 10^2), \quad \tau_\mu, \tau_{\log \sigma}, \phi_k \sim \text{half-Student-}t_3(0, 5), \quad \mu_k \sim \text{Beta}(2, 2).$$

*S2.3.2 Lab-Bayes model.* The Lab-Bayes model uses the Group model's posterior as an informative prior for a specific laboratory. If $(\hat{\mu}_0, \hat{\tau}_\mu, \widehat{\log\sigma_0}, \hat{\tau}_{\log\sigma}, \hat{\mu}_k, \hat{\phi}_k)$ are Group-level posterior summaries, then for that lab:

$$\mu_{\text{lab}} \sim \mathcal{N}(\hat{\mu}_0, \hat{\tau}_\mu^2), \quad \log\sigma_{\text{lab}} \sim \mathcal{N}(\widehat{\log\sigma_0}, \hat{\tau}_{\log\sigma}^2), \quad k_{\text{lab}} \sim \text{Beta}(\hat{\mu}_k\hat{\phi}_k, (1-\hat{\mu}_k)\hat{\phi}_k).$$

The likelihood is the DL-aware mixture applied to that lab's raw direct or secondary data.

*S2.3.3 Lab-Vague model.* The Lab-Vague model uses weakly informative priors:

$$\mu_{\text{lab}} \sim \mathcal{N}(0, 100^2), \quad \log\sigma_{\text{lab}} \sim \mathcal{N}(0, (2.5)^2), \quad k_{\text{lab}} \sim \text{Beta}(0.5, 0.5),$$

with the same DL-aware likelihood as above. This allows lab-specific data to dominate while remaining numerically stable.

Stan [1] is used to obtain posterior draws $(\mu^{(t)}, \sigma^{(t)}, k^{(t)})$ for each transfer type and model family; these draws are then used in the case-level construction described in Sections 2.3 and 2.4.

*S2.4. Monte Carlo Construction of the Case-Level Likelihood Ratio*

The samples from the posterior distribution of the parameters $\theta$ are used to provide a Monte Carlo representation of the case-level LR. Concretely, for each posterior draw $\theta^{(t)}$, HaloGen evaluates the full case-level likelihood under $H_p$ and $H_d$ using the same posterior state, and forms a per-draw LR

$$\text{LR}^{(t)} = \frac{P(E \mid H_p, \theta^{(t)})}{P(E \mid H_d, \theta^{(t)})}.$$

The collection $\{\text{LR}^{(t)}\}$ is the Monte Carlo representation of the posterior-marginalised LR.

The "paired per-draw" construction is therefore not a separate inferential approach, but an operational Monte Carlo strategy used in HaloGen. The mathematical definition of the LR is the ratio of the likelihood of the evidence under the competing propositions. In practice, evaluating the direct-transfer and secondary-transfer likelihood components at the same Monte Carlo draw preserves draw-level coherence between shared or corresponding uncertain model components and reduces avoidable simulation noise. Unpaired Monte Carlo schemes could in principle be used, but they would introduce additional

Monte Carlo variability and would make the draw-wise LR summaries less directly interpretable.

Summary statistics (median, quantiles) are reported only as a presentation choice and do not define a different inferential method.

*S2.5. Detection-Limit Robustness and Diagnostics*

HaloGen includes additional robustness checks to verify that moderate misspecification of $DL$ does not unduly distort the LRs.

The software recomputes paired per-draw LRs under three detection-limit settings:

$$DL_{\text{low}} = \frac{DL}{2}, \qquad DL_{\text{base}} = DL, \qquad DL_{\text{high}} = 2DL,$$

and across three data configurations:

- the observed data (mix of detects and non-detects),
- a synthetic "zeros-only" scenario (all non-detects),
- a synthetic "positives-only" scenario (no non-detects).

For each configuration and model family (*Group*, *Lab–Bayes*, *Lab–Vague*), HaloGen compares:

- the median $\log_{10}(\text{LR})$,
- the 10–90% quantile band of $\log_{10}(\text{LR})$.

If changes exceed pre-specified thresholds, the lockdown diagnostics report a warning. These checks complement the main-case analysis in Sections 2.3–2.4 and provide assurance that the results are not unduly sensitive to modest uncertainty in the detection limit. More details are provided in [2], Supplement S4.

### S3. The HaloGen Simulation Framework

The HaloGen script implements a computational workflow that evaluates DNA evidence via likelihood ratios (LRs), allowing direct comparison between different statistical models (*Group*, *Lab–Bayes*, *Lab–Vague*). The core workflow (Fig. S1) proceeds through parameter estimation, scenario setup, casewise LR computation, and output writing, using function and object names that match the current Gemini v14_v6 engine.

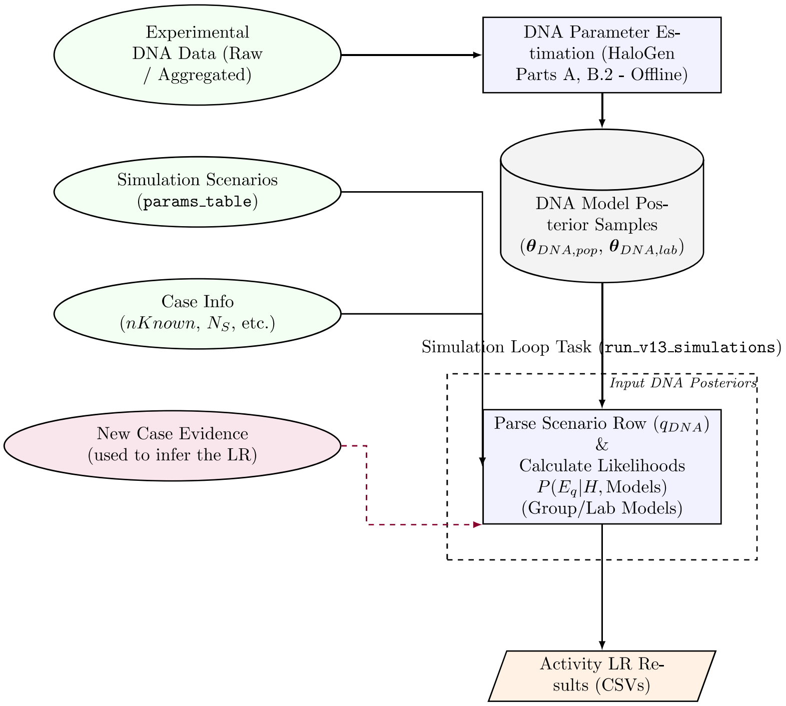


Figure S1: Conceptual workflow of the HaloGen simulation framework.

*Script-related definitions*

`group_post_samples_direct/secondary` Posterior samples for $(\mu, \log\sigma, k)$ (per transfer type) representing the cross-laboratory (Group) population distributions for direct and secondary transfer.

`post_lab_bayes_..._samples` Posterior samples for lab-specific parameters obtained using informative priors derived from the Group model (Lab–Bayes).

`post_lab_vague_..._samples` Posterior samples for lab-specific parameters obtained using weakly informative priors (Lab–Vague).

`stains_per_known/unknown` Integer vectors giving the number of replicate stains (DNA quantities) per known and per specified unknown contributor.

`params_table` Data frame where each row represents a simulation scenario or defined case. Contributor-specific stain quantities occupy columns such as `H1a, H1b, H2, ...`. Constructed via `setup_defined_case_params_table(...)` or `setup_simulation_scenario(...)`.

*(1) Inputs*

- *Experimental data.* Quantitative DNA results used to fit the Group and Lab models (Section 2.2). For the Group model, lab-wise summary statistics $(\hat{\mu}_l, \widehat{\log\sigma_l}, \hat{k}_l)$ are used; for Lab models, raw stain-level data per laboratory are used. Raw counts `fail_count` and `sample_size` are retained at the network level to support the empirical $F_0$ policy, which is constructed from pooled direct-transfer non-detection information stored in `direct_params`.

- *Simulation scenarios (`params_table`).* Defined cases or simulation scenarios specifying the observed DNA quantities for each known and unknown contributor across one or more stains.

- *Case information.* Numbers of specified knowns (`nKnown`), unknowns (`nUnknown`), actors (`nActors` $= N_S$), and the detection limit (`DL`).

*(2) Parameter estimation and priors (summary)*

The DL-aware, zero-inflated and censored lognormal parameters $\boldsymbol{\theta} = (\mu, \sigma, k)$ are estimated for each transfer type (direct, secondary) using Bayesian models implemented in Stan, as described in Section 2.2.

- *Group model.* A hierarchical cross-laboratory model provides posterior distributions for $(\mu_0, \tau_\mu, \log \sigma_0, \tau_{\log \sigma}, \mu_k, \phi_k)$, which in turn yield the population-level posterior samples `group_post_samples_direct` and `group_post_samples_secondary`.

- *Lab-specific models.* For each laboratory and transfer type, lab-specific DL-aware models are fitted with Group-level hyperparameters fixed to their posterior summaries

  to the raw stain-level data:

  - Lab–Bayes uses informative priors induced by the Group hyperposteriors.
  - Lab–Vague uses broad, weakly informative priors so that the lab's own data dominate the posterior.

  Full prior structures and hyperpriors are listed in Table S1 and are consistent with the estimation framework of Section 2.2.

All posterior draws (Group and Lab-specific) are saved as data frames of $(\mu, \log \sigma, k)$ and serve as direct inputs to the LR computation engine.

*S3.1. Choice of Priors for Bayesian Models*

This section summarises the prior distributions used in the HaloGen framework for modelling DNA transfer and activity-level hypotheses. The aim is to use priors that stabilise inference in small-sample settings without overwhelming the data, while remaining transparent and defensible for forensic application. We distinguish carefully between parameters governing experimental transfer processes within laboratories and the case-level parameters used in the exhaustive hypothesis set. All priors reported here correspond to those implemented in the current engine and are shared across the simulation studies presented in this paper.

Table S1: Priors used in HaloGen, as implemented in `Gemini_v14_v6`. Parameters are per lab and transfer mode unless labelled as hyperparameters.

| Model | Parameter | Prior (engine) | Rationale / notes |
|---|---|---|---|
| ***Group model: hierarchical partial pooling across laboratories*** | | | |
| | $\mu_{0,\mathrm{dir}}$; $\tau_{\mu,\mathrm{dir}}$ | $\mu_{0,\mathrm{dir}} \sim \mathcal{N}(0, 10^2)$<br>$\tau_{\mu,\mathrm{dir}} \sim$ Student-$t(3, 0, 5)$, constrained $> 0$ | Weakly informative global mean and shrinkage prior for laboratory-to-laboratory variability in direct transfer. |
| | $\mu_{0,\mathrm{sec}}$; $\tau_{\mu,\mathrm{sec}}$ | $\mu_{0,\mathrm{sec}} \sim \mathcal{N}(-2.2, 10^2)$<br>$\tau_{\mu,\mathrm{sec}} \sim$ Student-$t(3, 0, 5)$, constrained $> 0$ | Offset reflects typically lower secondary-transfer yields on the log scale. |
| | $\log \sigma_{0,\mathrm{dir}}$; $\tau_{\log\sigma,\mathrm{dir}}$ | $\log \sigma_{0,\mathrm{dir}} \sim \mathcal{N}(0, 10^2)$<br>$\tau_{\log\sigma,\mathrm{dir}} \sim$ Student-$t(3, 0, 5)$, constrained $> 0$ | Log-scale parameterisation ensures $\sigma > 0$; the Student-$t$ scale prior is stable and weakly informative. |
| | $\log \sigma_{0,\mathrm{sec}}$; $\tau_{\log\sigma,\mathrm{sec}}$ | $\log \sigma_{0,\mathrm{sec}} \sim \mathcal{N}(0.5, 10^2)$<br>$\tau_{\log\sigma,\mathrm{sec}} \sim$ Student-$t(3, 0, 5)$, constrained $> 0$ | Secondary-transfer offset reflects typically broader dispersion. |
| | $\mu_{k,\mathrm{dir}}$; $\phi_{k,\mathrm{dir}}$; $k_{\mathrm{lab,dir}}$ | $\mu_{k,\mathrm{dir}} \sim \mathrm{Beta}(2, 2)$<br>$\phi_{k,\mathrm{dir}} \sim$ Student-$t(3, 0, 5)$, constrained $> 0$<br>$k_{\mathrm{lab,dir}} \sim \mathrm{Beta}(\mu_k\phi_k,\ (1-\mu_k)\phi_k)$ | Mean–concentration hierarchy for the non-detect component $k$, with partial pooling across laboratories. |
| | $\mu_{k,\mathrm{sec}}$; $\phi_{k,\mathrm{sec}}$; $k_{\mathrm{lab,sec}}$ | $\mu_{k,\mathrm{sec}} \sim \mathrm{Beta}(2, 2)$<br>$\phi_{k,\mathrm{sec}} \sim$ Student-$t(3, 0, 5)$, constrained $> 0$<br>$k_{\mathrm{lab,sec}} \sim \mathrm{Beta}(\mu_k\phi_k,\ (1-\mu_k)\phi_k)$ | Same mean–concentration hierarchy for the secondary-transfer model. |
| ***Lab–Bayes model: lab-specific model with Group-informed hyperparameters*** | | | |
| | $\mu_0$; $\tau_\mu$; $\log\sigma_0$; $\tau_{\log\sigma}$; $\mu_k$; $\phi_k$ | Fixed at Group posterior summaries, separately by transfer type. | Implements partial pooling by borrowing strength from the Group model while fitting each laboratory to its own data. |
| | $\mu_{\mathrm{lab}}$; $\log\sigma_{\mathrm{lab}}$; $k_{\mathrm{lab}}$ | Same non-centred lab-specific model as below, using the fixed Group-informed hyperparameters. | The engine fits each laboratory with a non-centred parameterisation for sampler stability. |
| ***Lab–Vague model: standalone lab-specific model with weak constraints*** | | | |
| | $\mu_0$; $\tau_\mu$ | $\mu_0 = 0$; $\tau_\mu = 10$ | Broad location prior on the log-quantity scale. |
| | $\log\sigma_0$; $\tau_{\log\sigma}$ | $\log\sigma_0 = 0$; $\tau_{\log\sigma} = 5$ | Broad dispersion prior without heavy tails. |
| | $\mu_k$; $\phi_k$; $k_{\mathrm{lab}}$ | $\mu_k = 0.5$; $\phi_k = 1$, so that $k_{\mathrm{lab}} \sim \mathrm{Beta}(0.5, 0.5)$. | Deliberately weak prior for $k$; permits extreme values when laboratory data are sparse because no shrinkage is used. |
| ***Case-level policy: exhaustive hypothesis set*** | | | |
| | $F_0$ | Empirical Jeffreys-smoothed Beta distribution with clamping and an extra-cautious floor. | Distinct from $k$; governs the non-detection probability for a relevant unobserved actor in the exhaustive hypothesis set. |

*Notes on choices.* (i) For Group-level scale parameters we use half-Student-$t$ priors (implemented as student_t$(3, 0, 5)$ with positive constraints in Stan), which are lighter-tailed than half-Cauchy and empirically stable in this application. (ii) In the current engine implementation, the standalone Lab–Vague model uses a Jeffreys-type prior for the experimental non-detect component $k$, namely $k \sim \text{Beta}(0.5, 0.5)$ (arising from $\mu_k = 0.5$, $\phi_k = 1$ in the mean–concentration parameterisation). This choice is intentionally weakly informative and allows substantial mass near 0 and 1 when laboratory data are sparse. (iii) The case-level fail-rate term $F_0$ is conceptually distinct from the experimental parameter $k$: $k$ governs non-detects within the transfer likelihood, whereas $F_0$ governs the probability that an (unobserved) actor leaves no detectable DNA under the activity-level hypothesis set. The engine applies an empirical Jeffreys-smoothed update for $F_0$ with clamping and an extra-cautious floor.

*Sensitivity and reporting.* Likelihood ratios are computed using paired posterior draws and summarised as medians with 10-90% intervals on the $\log_{10}$ scale. Where reported, prior-sensitivity checks for the Lab-Vague $k$ component (e.g. $\text{Beta}(1, 1)$ or $\text{Beta}(3, 3)$ in place of $\text{Beta}(0.5, 0.5)$) primarily affect tail behaviour and interval width rather than the median. A limitations note is flagged if the $\log_{10} \text{LR}_{\text{none}}$ interval exceeds three units or if the effective number of finite paired draws is low.

*(3) Casewise simulation loop (`run_v14_simulations`)*

For each row of `params_table`, the core engine proceeds as follows:

- *Parse contributor-wise quantities.* The function extracts the stain quantities for all specified contributors and builds the internal representation required by the exhaustive framework (Section 2.4), including `stains_per_known`, `stains_per_unknown` and the actor-count $N_S$.
- *Per-draw DL-aware likelihoods.* For each Monte Carlo draw $t$ from the relevant posterior(s), the engine evaluates per-stain likelihoods under the censored mixture model of Section 2.1. The base (unconditional) per-observation likelihood used in model fitting is

$$L(q \leq DL \mid \theta, DL) = k + (1-k)\,\Phi(z_0), \qquad L(q > DL \mid \theta, DL) = (1-k)\, f_{\text{LN}}(q; \mu, \sigma),$$

where $z_0 = (\log DL - \mu)/\sigma$.

At the *case-evaluation* stage, when assembling contributor-wise case-level likelihoods $t_c$ and $s_c$ for detected quantities $q > DL$, HaloGen applies the conditioning rule of Section 2.3: for each posterior draw,

$$L_D^{\text{cond}}(q) = \frac{f_{\text{LN}}(q; \mu_D, \sigma_D)}{1 - \Phi\big((\log DL - \mu_D)/\sigma_D\big)}, \qquad L_S^{\text{cond}}(q) = \frac{f_{\text{LN}}(q; \mu_S, \sigma_S)}{1 - \Phi\big((\log DL - \mu_S)/\sigma_S\big)}.$$

Both direct and secondary transfer likelihoods are conditioned to the detected domain $[DL, \infty)$, so that contributor-wise case-level densities are evaluated conditional on detection , and the mixture-model terms involving $k$ and the sub-threshold mass are removed at case level. Non-detects ($q \leq DL$) are not assigned contributor-wise case-level densities; their impact is captured via the actor fail-rate $F_0$ as described in Section 2.3.1.

- *Case-level fail-rate injection $F_{0,t}$.* For each draw $t$, the engine constructs a case-level fail rate $F_{0,t}$ using the empirical–Jeffreys policy with clamp and tail floor (Section 2.3.1):

$$\begin{aligned}
F_{0,t} &\sim \text{Beta}(f + 0.5,\ n - f + 0.5), \qquad\qquad J = \frac{f + 0.5}{n + 1}, \\
F_{0,t}^{\text{policy}} &= \min\{\max(F_{0,t}, J),\ 1 - J\}, \\
F_{0,t}^{\text{tail}} &= k_D^{(t)} + (1 - k_D^{(t)})\,\Phi\big((\log DL - \mu_D^{(t)})/\sigma_D^{(t)}\big), \\
F_{0,t} &= \max\{F_{0,t}^{\text{policy}}, F_{0,t}^{\text{tail}}\}.
\end{aligned}$$

  In an elemental hypothesis with $x$ specified actors (out of $N_S$), the unobserved-actor contribution is $\big(F_{0,t}\big)^{(N_S - x)}$.

- *Exhaustive LR computation.* For each model (*Group*, *Lab–Bayes*, *Lab–Vague*), the function `compute_caseBF_stainwise_one_model_PM(...)` assembles per-draw case Bayes factors by summing over all elemental hypotheses $H_{\mathcal{O}}$ under the exhaustive framework, using a *paired per-draw* construction: numerator and denominator always share the same draw index $t$ (Section 2.4). The "PM" wrappers enforce the global policy options

```
options(
  gemini.case_bf_mode      = "condition_on_scene_open",
  gemini.f0.policy         = "empirical_beta_clamped",
  gemini.f0.autoinject     = TRUE,
  gemini.f0.extra_cautious = TRUE
)
```

so that all models share the same $F_0$ treatment; they differ only in their $(\mu, \sigma, k)$ posterior sources.

*(4) Outputs and writer integration*

Each call to `run_v14_simulations` returns triads (median, 10%, 90%) of $\log_{10} \mathrm{BF}_{\mathrm{case}}$ for each model. The Batch3 / Batch_v16 writers additionally call:

- `compute_contributor_CASELR_quantiles_one_model_PM(...)` to obtain contributor-specific CASE-LRs; and
- `compute_LRnone_quantiles_one_model_PM(...)` to obtain $\mathrm{LR}_{\mathrm{none}}$.

Results are written to the workbook sheets *Consolidated_Log10_BF_Case*, *Audit_Numeric*, *LR_none*, and *Policy*.

*Policy integrity and sourcing.* To guarantee that the case-level detect rule and case-level $F_0$ policy are applied consistently, the policy is set *before* sourcing the engine and writer and checked *after* sourcing:

```
OriginalF0Policy_On()
assign("raw_data_all_labs", RAW_ENGINE, envir = .GlobalEnv)
assign("detection_limit_raw", DETECTION_LIMIT, envir = .GlobalEnv)

sys.source(ENGINE_FILE, envir = .GlobalEnv)
sys.source(BATCH3_FILE, envir = .GlobalEnv)
OriginalF0Policy_Assert()
```

A guard ensures that `direct_params` and the baseline $J$ are computed from the full network (all labs); if not, the files are re-sourced with `lab_id_to_analyze` cleared.

*Important.* Analysts should not recompute LRs outside the engine. All numeric results (case BF, contributor CASE-LR, and $\mathrm{LR}_{\mathrm{none}}$) are to be obtained via the engine entry points, such as

```
compute_caseBF_stainwise_one_model(
  <model_key>,
  data.frame(H1 = q),
  nKnown = 1, nUnknown = 0,
  DL = DL
)
```

which preserves the case-level detect rule, the exhaustive summation over elemental hypotheses, and the enforced $F_0$ policy.

*(5) Posterior diagnostics, draws, and convergence*

All Bayesian models in HaloGen are fitted using Stan's dynamic Hamiltonian Monte Carlo sampler. Because posterior draws feed directly into the per-draw likelihood calculations, draw quality, convergence, and effective sample size are critical.

*Number of draws used by HaloGen.* For each transfer type (direct, secondary) and for each model (*Group*, *Lab–Bayes*, *Lab–Vague*), HaloGen uses a unified Stan configuration:

- 8 parallel chains, each with 8,000 iterations (4,000 warm-up and 4,000 post–warm-up);
- yielding exactly 32,000 post–warm-up draws per model and per transfer type.

No thinning is applied. Unless the user explicitly requests downsampling, the LR engine consumes all 32,000 draws for each model. Each Monte Carlo index $t$ provides a paired realisation of the direct and secondary parameters in the per-draw LR construction (Section 2.4), ensuring that cross-arm parameter uncertainty is handled coherently. Effective draw counts for all LR summaries are reported by the software. Unless otherwise specified, HaloGen uses a unified Stan configuration:

*Convergence and HMC diagnostics.* For every fitted model, the following diagnostics are monitored:

- the potential scale reduction statistic $\widehat{R}$ for all scalar parameters (including $\mu$, $\log \sigma$, $k$, and all hierarchical hyperparameters), with a requirement that $\widehat{R} \leq 1.01$;
- bulk and tail effective sample sizes (ESS), with a target of several hundred effective draws per parameter and substantially higher ESS for $(\mu, \log \sigma, k)$ in the regions that drive the likelihood;
- zero divergent transitions after warm-up (target: no divergences);

- tree-depth and energy-based diagnostics (e.g. E-BFMI) to detect geometries that may cause bias or slow mixing.

To achieve stable geometry for heavy-tailed likelihood regions and censoring-driven curvature, the sampler uses a high acceptance threshold (`adapt_delta = 0.995`) and a maximum tree depth of 15. In particularly sparse laboratories, single-chain fits are permitted provided trace plots, autocorrelation functions, and ESS demonstrate stable and well-mixed posteriors.

*Posterior predictive checks.* Posterior predictive simulations are routinely used to assess model fit. For each converged model, replicated datasets are drawn from the DL-aware mixture, and we verify that:

- the simulated distribution of positive quantities spans the observed range;
- the simulated non-detect rate matches the empirical non-detect rate (within posterior uncertainty);
- the implied dropout behaviour—combining 'no-transfer events' ($k$) and sub-threshold mass $\Phi((\log DL - \mu)/\sigma)$—remains physically plausible for both direct and secondary transfer.

*Propagation of uncertainty.* HaloGen never substitutes point estimates of $(\mu, \sigma, k)$. Instead, each Monte Carlo index $t$ provides:

- a full parameter triplet $(\mu_t, \sigma_t, k_t)$ for both direct and secondary transfer, and
- a corresponding dropout mass $F_{0,t}$ under the empirical-$F_0$ policy (with extra-cautious clamping).

These draws are applied consistently in both numerator and denominator of the likelihood ratio. This guarantees that:

- parameter uncertainty, cross-laboratory variation, lab-specific deviations, and dropout uncertainty are fully propagated; and
- all evidential likelihood ratios and Bayes factors are evaluated via Monte Carlo marginalisation over the posterior is used, rather than plug-in point estimates for the model parameters. Posterior marginalisation is performed at the level of the complete case-level likelihood (and hence the LR), not by integrating each primitive likelihood component separately.

Simulations with very wide 10–90 % intervals for $\log_{10}$ LR (e.g. spanning more than three orders of magnitude) or with low ESS on key parameters are flagged for caution and may trigger additional MCMC runs or targeted sensitivity analyses.

### S4. Generalisation of the Effect of Increased Unknowns

This supplement provides a simplified mathematical illustration of how the presence of multiple specified unknown contributors can influence the likelihood ratio (LR) for a Person of Interest (POI) when conditioning on a single actor ($N_S = 1$).

> **Important note.** This section uses a simplified analytic model intended solely to illustrate mathematical trends. For clarity:
>
> - Posterior–predictive likelihoods $(t_1, s_1, \tau', s')$ are treated as fixed scalars.
> - Direct/secondary likelihoods are assumed unconditional (the asymmetric detect rule of the case model is *not* applied).
> - All specified unknown contributors are assumed to have identical likelihoods.
> - The case-level fail rate $F_0$ is treated as a constant (the operational model uses per-draw $F_{0,t}$).
>
> HaloGen's operational LR always uses the full exhaustive-propositions framework described in the main Methods.

*Notation*

Let the POI be contributor 1, and let there be $u$ specified unknown contributors indexed $j = 2, \ldots, u+1$. For each contributor we use posterior–predictive likelihoods:

$$
\begin{aligned}
t_1 &= P(E_1 \mid \text{direct (POI)}),\\
s_1 &= P(E_1 \mid \text{secondary (POI)}),\\
\tau' &= P(E_j \mid \text{direct (unknown } j)),\\
s' &= P(E_j \mid \text{secondary (unknown } j)),\\
F_0 &= P(\text{unobserved actor leaves no detectable DNA}).
\end{aligned}
$$

*Elemental Probabilities for $N_S = 1$*

*(1) $H_p$: The POI is the actor.* The POI's DNA is from direct transfer; all $u$ specified unknowns contribute innocently:

$$P(E \mid H_p) = t_1 (s')^u.$$

*(2) $H_{d,j}$: Specified unknown $j$ is the actor.* Unknown $j$'s DNA arises via direct transfer; the POI and the other unknowns transfer innocently:

$$P(E \mid H_{d,j}) = \tau' \, s_1 \, (s')^{u-1}.$$

*(3) $H_{d,UO}$: An unobserved actor is the sole actor.* All specified individuals contribute innocently and the single actor is unobserved:

$$P(E \mid H_{d,UO}) = s_1 (s')^u F_0.$$

*Likelihood Ratio*

We first consider the case $u \geq 1$, where $u$ denotes the number of specified unknown contributors available as alternative direct-transfer actors under the defence proposition. The LR comparing $H_p$ with the set of defence hypotheses, namely one of the $u$ specified unknowns or one unobserved actor, is

$$LR_{POI} = \frac{P(E \mid H_p)}{\displaystyle\sum_{j=2}^{u+1} P(E \mid H_{d,j}) + P(E \mid H_{d,UO})}.$$

Substituting the expressions above gives

$$LR_{POI} = \frac{t_1 (s')^u}{u \left\{ \tau' s_1 (s')^{u-1} \right\} + s_1 (s')^u F_0}.$$

For $u \geq 1$, cancelling $(s')^{u-1}$ gives

$$LR_{POI}(u \geq 1) = \frac{t_1 \, s'}{s_1 \, (u\tau' + s' F_0)}. \quad \text{(S1)}$$

*Limiting case with no specified unknown contributors.* The preceding expression assumes $u \geq 1$, because the denominator contains a sum over specified unknown contributors and the algebra cancels $(s')^{u-1}$. When $u = 0$, there are no specified unknown contributors to act as alternative detected direct-transfer actors. The defence set therefore contains only the unobserved-actor alternative. In that case,

$$P(E \mid H_p) = t_1, \qquad P(E \mid H_{d,UO}) = s_1 F_0,$$

and hence

$$LR_{POI}(u = 0) = \frac{t_1}{s_1 F_0}. \tag{S2}$$

Thus, the $u = 0$ case is a separate limiting case, not a direct substitution into Eq. (S1).

*Behaviour as the Number of Specified Unknowns Increases*

For $u \geq 1$, if the unobserved-actor term $s' F_0$ is small relative to $u\,\tau'$, then Eq. (S1) gives

$$LR_{POI}(u) \approx \frac{t_1 s'}{s_1(u\tau')} = \frac{C}{u}, \qquad C = \frac{t_1 s'}{s_1 \tau'}.$$

Thus, under these simplifying assumptions and for $u \geq 1$,

$$LR_{POI}(u) \approx \frac{1}{u} \times C.$$

**Interpretation.** With more specified unknown contributors available as plausible alternative direct-transfer actors, the denominator of the LR increases approximately linearly in $u$. Hence, under the simplifying assumptions above, the LR for the POI decreases approximately in proportion to $1/u$.

**Relation to the main Methods.** This simplified behaviour is consistent with the intuition embodied in the full exhaustive-propositions model: adding more plausible specified contributors redistributes evidential weight across a larger set of defence alternatives. The special case $u = 0$ is handled separately because no specified unknown alternatives are present.

### References

[1] B. Carpenter, A. Gelman, M. D. Hoffman, D. Lee, B. Goodrich, M. Betancourt, M. Brubaker, J. Guo, P. Li, A. Riddell, Stan: A Probabilistic Programming Language, Journal of Statistical Software 76 (1) (2017) 1–32.

[2] P. Gill, Ø. Bleka, Likelihood Ratios Given Activity-Level Propositions for DNA Transfer Evidence: Practical Implementation and Simulation Studies Using the HaloGen Engine (Part II), bioRxiv (2026). doi:10.64898/2026.02.06.703509.
URL https://www.biorxiv.org/content/10.64898/2026.02.06.703509v1
